# A mitochondria-specific mutational signature of aging: increased rate of A>G substitutions on a heavy chain

**DOI:** 10.1101/2021.12.03.460832

**Authors:** A. G. Mikhaylova, A. A. Mikhailova, K. Ushakova, E.O. Tretiakov, D. Iliushchenko, V. Shamansky, A. Iurchenko, M. Zazhytska, E. Kozenkova, E. Zdobnov, V. Makeev, V. Yurov, M. Tanaka, I. Gostimskaya, Z. Fleischmann, S. Annis, M. Franco, K. Wasko, W.S Kunz, D.A. Knorre, I. Mazunin, S. Nikolaev, J. Fellay, A. Reymond, K. Khrapko, K. Gunbin, K. Popadin

## Abstract

The mutational spectrum of the mitochondrial DNA (mtDNA) does not resemble any of the known mutational signatures of the nuclear genome and variation in mtDNA mutational spectra between different organisms is still incomprehensible. Since mitochondria is tightly involved in aerobic energy production, it is expected that mtDNA mutational spectra is affected by the oxidative damage. Assuming that oxidative damage increases with age, we analyze mtDNA mutagenesis of different species. Analysing (i) dozens thousands of somatic mtDNA mutations in samples of different age (ii) 70053 polymorphic synonymous mtDNA substitutions, reconstructed in 424 mammalian species with different generation length and (iii) synonymous nucleotide content of 650 complete mitochondrial genomes of mammalian species we observed that the frequency of A_H_>G_H_ substitutions (_H_ - heavy chain notation) is twice higher in species with high versus low generation length making their mtDNA more A_H_ poor and G_H_ rich. Considering that A_H_>G_H_ substitutions are also sensitive to the time spent single stranded (TSSS) during asynchroniuos mtDNA replication we demonstrated that A_H_>G_H_ substitution rate is a function of both species-specific generation length and position specific TSSS. We propose that A_H_>G_H_ is a mitochondria-specific signature of oxidative damage associated with both aging and TSSS.

## Introduction

Molecular evolution is a function of boht mutagenesis and selection. In order to uncover selection forces, it is crucial to reconstruct the mutational process. It has been suggested for example, that an excess of G nucleotides in the mitochondrial genome (mtDNA) of long-lived mammals is a result of selection, favoring more stable genomes in long-lived species (Lehmann et al. 2006). However, this conclusion can be premature without a comparison of mutational processes between short- and long-lived mammals. Indeed, significant changes in mtDNA mutational spectra between different species have been shown (Belle et al. 2005; Montooth and Rand 2008), however, no driving factors explaining this variation has been proposed till now.

Interestingly, a similar gap of knowledge on mtDNA mutational spectra exists on the comparative-tissues level. Pan-cancer studies have shown that mtDNA has a unique mutational signature that differs from all known nuclear signatures (Yuan et al. 2017; Ju et al. 2014). Moreover, well known strong exogenous mutagens such as tobacco smoke in lung cancers of smokers or ultraviolet light in melanomas do not show expected effects on the mitochondrial mutational spectrum (Yuan et al. 2017; Ju et al. 2014). Thus, the main mutagen of mtDNA as well as the causes of variation in mtDNA mutational spectra are unknown on both comparative-tissues and comparative-species scales.

The widely accepted expectation is that mitochondrially produced reactive oxygen species (ROS) can damage mtDNA (Alexeyev 2009). The well-documented ROS-induced signature is the modification of the guanine (G) DNA base to 7,8-Dihydro-8-oxo-20-deoxyguanosine (8-oxodG), which after mispairing with adenine, leads to G > T transversion mutations. Although G>T substitutions are considered to be the hallmark of oxidative damage in the nuclear DNA (COSMIC signature 18) (Fraga et al. 1990; Alexandrov et al. 2013; Kucab et al. 2019), it is rather rare in mtDNA (Yuan et al. 2020) and doesn’t significantly increase with age in mtDNA (Kennedy et al. 2013) (Yuan et al. 2020)(Guliaeva, Kuznetsova, and Gaziev 2006). So, up to now, there is no well-established mutational signature of oxidative damage in mtDNA (Zsurka et al. 2018).

Taking into account recent progress in deciphering the variation in mutational spectra of the nuclear genome as a function of different cancer types (G. Koh et al. 2021), environmental agents (Kucab et al. 2019), gene knockouts (Zou et al. 2021), human populations (Harris and Pritchard 2017) and primate species (Moorjani et al. 2016), here we focus on mtDNA and perform a large-scale reconstruction of its mutational spectra across hundreds of mammalian species. Considering the tight association of the level of mtDNA metabolism (and thus potential mtDNA mutagens) with species-specific life-history traits, we aimed to uncover a correlation between the mtDNA mutational spectrum and life-history traits. Using collections of (i) somatic mtDNA mutations in mice and humans, (ii) polymorphic synonymous substitutions in hundreds of mammalian species, and (iii) nucleotide content in whole mitochondrial genomes of mammals, we observed one universal trend: A_H_>G_H_ substitutions (_H_ - heavy strand notation, see methods) positively correlates with the generation length. Considering an additional association of the A_H_>G_H_ substitutions with the time spent single stranded (TSSS) during asynchronous mtDNA replication and numerous literature data about A>G substitutions, we propose that the increased A_H_>G_H_ in mtDNA of long-lived mammals is a mutational signature of an age-associated oxidative damage, specific to single-stranded DNA. The described variation in the mtDNA mutational spectrum should be taken into account in somatic, population and evolutionary analyses of mtDNA.

## Results

### (1) Frequency of de novo A_H_>G_H_ mutations increases with age in soma and germ-line

Mitochondrial genome is characterized by the strong strand asymmetry in the nucleotide content: the heavy strand (H-strand) is guanine rich (G_H_) and cytosine poor (C_H_), while the light strand (L-strand) is opposite: cytosine rich (C_L_) and guanine poor (G_L_). The mutagenic explanation of this asymmetry is based on an assumption that mtDNA heavy strand, being single stranded during asynchronous replication, is more susceptible to two of the most common mutations in mtDNA: C_H_>T_H_ and A_H_>G_H_ leading to a deficit of C_H_ and an excess of G_H_. Analyses of complete mitochondrial genomes of mammals showed additionally that this nucleotide asymmetry forms a gradient along mtDNA (Reyes et al. 1998; Tanaka and Ozawa 1994; Faith and Pollock 2003): a global deficit of C_H_ over T_H_ and A_H_ over and G_H_ at the third codon positions is becoming more pronounced along the major arc from COX1 to CYTB (Figure 1A). This gradient also supports the asynchronous mode of mtDNA replication and a mutagenic effect of the time spent single-stranded (TSSS): the two most common transitions C_H_>T_H_ and A_H_>G_H_ are more frequent in the region of CYTB which spent significantly more time in a single stranded state as compared to COX1 (Raina et al. 2005) (Figure 1A). Recently, a large collection of somatic mutations, generated by a highly sensitive duplex sequencing approach, allowed to precisely reconstruct the C_H_>T_H_ and A_H_>G_H_ gradients and to unambiguously confirm the mutagenic effect of TSSS during the asynchronous replication (Figure 1A) (Sanchez-Contreras et al. 2021).

**FIGURE 1:**
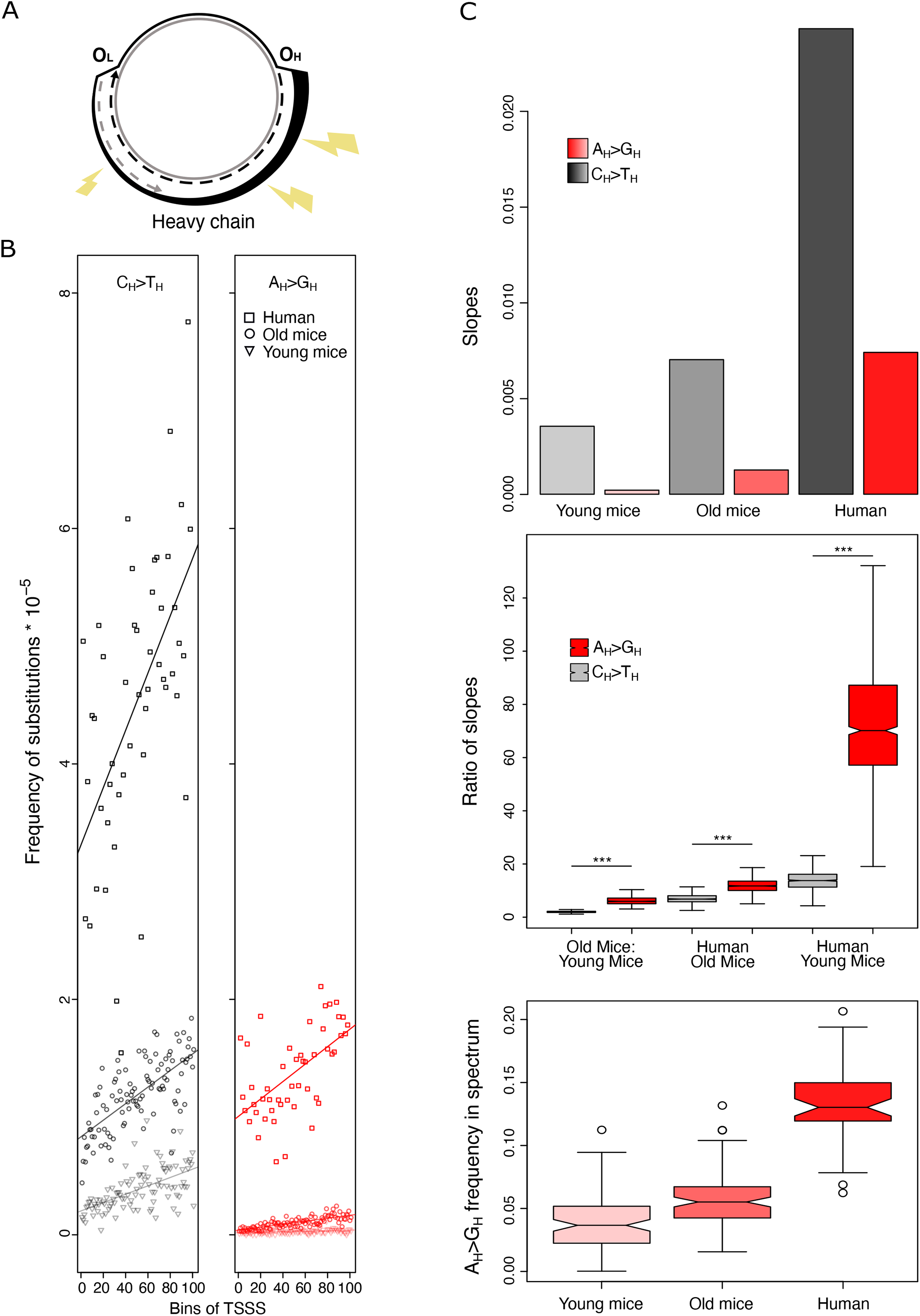
A_H_>G_H_ mtDNA mutational gradient is increasing with the sample age. 1A. Asynchronous replication of mtDNA is associated with a long time spent single stranded (TSSS) by the parental heavy chain. TSSS in turn is associated with the high frequency of two the most common mtDNA transitions: C_H_>T_H_ and A_H_>G_H_. daughter heavy chain - dashed black line; parental heavy chain - bold black line, with bolndess reflecting the TSSS; daughter light chain - dashed gray line; parental light chain - solid gray line. OH - origin of replication of daughter heavy chain, OL - origin of replication of daughter light chain. 1B. Gradients of C_H_>T_H_ and A_H_>G_H_ mutations along the major arc of mtDNA are more pronounced in humans versus old mice and in old mice versus young mice. Both intercepts and slopes are increasing with sample age (see supplementary materials 1.1). 1C. A_H_>G_H_ substitution rate is increasing faster in aged samples (see also supplementary materials 1.1) upper panel: barplots visualize the slopes of the linear regressions (see fig 1B) between the mutation frequency and TSSS. middle panel: A_H_>G_H_ slopes increase faster with age as compared to C_H_>T_H_ slopes. Boxplots are based on the ratio of slopes derived from 1000 bootstrapped samples. bottom panel: frequency of A_H_>G_H_ in the total mutational spectrum is increasing with age (p-values from all three pairwise comparisons are less than 1.583e-08, Mann-Whitney U test). ‘***’ marks p values < 0.001.

Recent confirmation of the mutational nature of the mtDNA gradients (Sanchez-Contreras et al. 2021) provides a solid ground for further investigations of the mtDNA mutational spectra. It was noted for example, that the positive G_H_/A_H_ gradient significantly differs between primate species being higher in species with longer gestation time, while T_H_/C_H_ gradient didn’t show strong species-specific variations (Raina et al. 2005). This suggests that the A_H_>G_H_ mutations, shaping the G_H_/A_H_ gradient, can be sensitive to some mutagens associated with gestation time or other life-history traits. Due to existence of a positive correlations between gestation time, body size, longevity, which in turn are associated with generation length and the level of mitochondrial metabolism we can expect a differences in mtDNA mutagenesis between species with different life-hisotry traits. To test this hypothesis we analyzed a dataset of somatic mtDNA mutations of mice and men.

To test a potential sensitivity of the two most common transitions to life-history traits we reanalysed the recent dataset of somatic de novo mtDNA mutations, obtained by the duplex sequencing approach (Sanchez-Contreras et al. 2021). Sanchez-Contreras et al had three groups of samples: young mice (4–5 months), old mice (26 months) and humans (10-90 years), which allow us to test an effect of the age on the mtDNA mutational spectrum. First of all, for comparative purposes, we plotted C_H_>T_H_ and A_H_>G_H_ gradients on the same scale (Figure 1B) and rerun the linear regression models describing the frequency of mutations as a function of TSSS (Supplementary Materials 1.1, Methods). As expected, and as has been shown in the paper (Sanchez-Contreras et al. 2021), we observed that both gradients demonstrate increased intercepts and slopes with age (Figure 1B, Supplementary Materials 1.1). Second, we focused on the slopes of the linear regressions as a proxy of a sensitivity of mutations to TSSS. We compared the relative increase of slopes from young mice, to old mice, to humans (Figure 1C upper panel, Supplementary Materials 1.1) and observed that the rate of increase of slopes is higher for A_H_>G_H_ as compared to C_H_>T_H_: it is 6.05 (1.98 for C_H_>T_H_) fold higher in old versus young mice, 11.7 (6.9 for C_H_>T_H_) fold higher in humans versus old mice and 70.86 (13.7 for C_H_>T_H_) fold higher in humans versus young mice. 1000 bootstrap resamplings with consequent recalculation of six slopes confirmed the robustness of this result: the increase in A_H_>G_H_ slope with age is higher than the increase in C_H_>T_H_ slope (Figure 1C middle panel, all p-values < 10^−16^, Mann-Whitney U test). A_H_>G_H_ mutations, due to the faster increase in slope with age, are expected to contribute proportionally more to the aged samples as compared to C_H_>T_H_. We estimated the total mutational spectrum of young mice, old mice and humans and observed that the total fraction of A_H_>G_H_ is indeed increasing with age (Figure 1C lower panel), while the fraction of C_H_>T_H_ shows no monotonic increase with age (Supplementary Materials 1.1). Altogether, using the datasets of somatic mtDNA mutations, obtained by a highly sensitive duplex sequencing approach, we uncovered that A_H_>G_H_ is more sensitive to age as compared to C_H_>T_H_.

Assuming a similarity of mtDNA mutagenesis in somatic and germ-line tissues we expect to observe also an excess of A_H_>G_H_ in aged germ-line tissues. Indeed, recent deep sequencing of de novo mtDNA mutations in aged versus young mice oocytes confirmed that the strongest hallmark of oocyte aging is an increased fraction of A_H_>G_H_ substitutions (Arbeithuber et al. 2020).

Additionally we performed an analysis of the human de novo mtDNA mutations as a function of the female reproductive age, which is a proxy for oocyte age. It has been shown that the number of de novo mtDNA mutations in children is increasing with maternal age (Rebolledo-Jaramillo et al. 2014; Wei et al. 2019), however, no age-related changes in mtDNA mutational spectra have been documented yet due to low sample size of the de novo mutations. We reanalyzed all of the de novo germline mutations from two recent studies of mother-offspring pairs (Rebolledo-Jaramillo et al. 2014; Wei et al. 2019). Due to low available sample sizes all the analyses of de novo mutations in mother-offspring pairs were only suggestive but they onsistently showed a trend of increased fraction of A_H_>G_H_ with oocyte age (Supplementary Materials 1.2). Thus our results suggest the frequency of de novo A_H_>G_H_ mutations increases with age in both soma and germ-line of mammals.

### (2) A_H_>G_H_s are more prevalent in mammals with high generation length: an evidence from polymorphism-derived neutral mutational spectra

The variation in mtDNA mutational spectra between different species (Montooth and Rand 2008; Belle et al. 2005) has no overarching explanation till now. Our findings (Fig 1) and literature data (Arbeithuber et al. 2020) suggest that this variation, and particularly a fraction of mtDNA A_H_>G_H_ transitions, can be associated with aging. Thus we hypothesize that species-specific mtDNA mutational spectra depends on the generation length which is in turn a good proxy of oocyte age in mammals. Because mammalian oocytes are arrested from birth to puberty, which takes weeks (for mice) or decades (for humans) (Von Stetina and Orr-Weaver 2011) we can use the species-specific generation length as a natural proxy for oocyte longevity in different mammalian species. Additionally, because oocytes are the only lineage through which mtDNA is transmitted (with rare exceptions of paternal inheritance) from generation to generation in mammals (Sato and Sato 2017) we expect to observe a correlation between the species-specific mtDNA properties and the generation length (a proxy for oocyte longevity) in different mammalian species. The generation length is defined as “the average age of parents of the current cohort” (Pacifici et al. 2013; Tacutu et al. 2013); it is available for the vast majority of mammalian species (Pacifici et al. 2013; Tacutu et al. 2013) and is also associated with numerous ecological (body mass, litter size, effective population size) and physiological (basal metabolic rate) parameters of mammalian species (Ollason 1987; Damuth 1987).

Numerous mitochondrial sequences coming from ecological, evolutionary, and population genetics studies of different species (Hebert et al. 2003) provide a valuable source of mtDNA polymorphisms, which based on our in-house pipeline (see Methods and Supplementary Materials 2.1) were used to reconstruct the mutational spectrum of mammalian species. Briefly, we downloaded all available nucleotide sequences of mitochondrial protein-coding genes of mammals, obtained multiple codon alignment for each gene of each species, rooted the mitochondrial within-species tree by the nearest neighbor sequence from another species, reconstructed the ancestral sequences in each inner node, obtained a list of polarized single-nucleotide substitutions and normalized yjrm by the frequency of ancestral nucleotides. Focusing on the most neutral 70,053 substitutions, located within the four-fold degenerate synonymous sites, we reconstructed the neutral mutational spectrum for 611 mammalian species (Supplementary Materials 2.2). The average mutational spectrum of all mammalian species (Figure 2A) demonstrates strong excess of C_H_>T_H_ and A_H_>G_H_ substitutions which have been shown in previous studies (Ju et al. 2014; Yuan et al. 2020).

**FIGURE 2:**
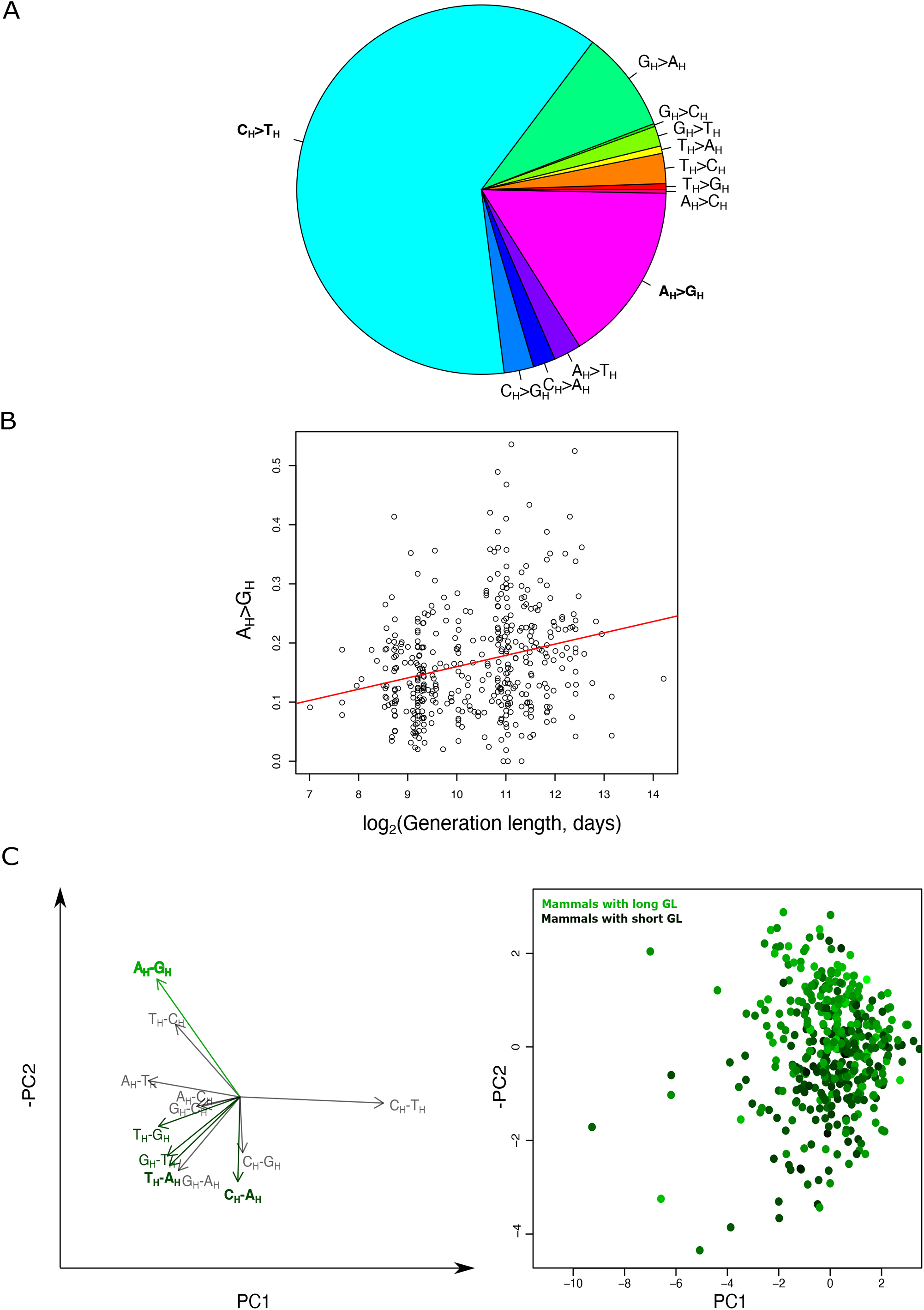
variation in neutral mtDNA mutational spectrum of mammals is driven by the generation length 2A. An average mtDNA mutational spectrum of mammalian species (N = 611). Mutational spectrum is a probability of each nucleotide to mutate to each other based on the observed and normalized frequencies of twelve types of nucleotide substitutions in four fold degenerate synonymous sites of all available within-species polymorphisms of mtDNA protein-coding genes. See also Supplementary Materials 2.2. 2B. Mutational spectra vary with species-specific generation length (N = 424). A_H_>G_H_ is the type of substitutions, frequency of which stronger correlated with the generation length. It shows approximately two fold difference between the mammalian with very short and very long generation length. See additionally Supplementary Materials 2.4. 2C. The principal component analysis (PCA) of mtDNA mutational spectra of mammalian species (N = 424). Left panel: the biplot of the principal component analyses (first and the second components explains 16% and 12% of variation correspondingly). C_H_>T_H_ has the highest loading on the first principal component while A_H_>G_H_ has the highest loading on the second principal component. Note that we plotted negative PC2 to make it positively correlated with generation length. Right panel: The second principal component correlates with the generation length in mammals. Generation length is color-coded from dark green (the shortest generation length) to light green (the longest generation length). See also Supplementary Materials 2.5.

To focus on the species-specific variation in mutational spectra we analysed in details the MT-CYB gene, which is (i) the most common in our database: it contained 56% of all extracted substitutions (39,112 out of 70,053 used to draw Figure 2A) and (ii) in all mammalian species it is located close to the origin of replication of the heavy chain (O_H_) and thus is characterized by a high TSSS (Figure 1). The comparison of the *MT-CYB*-derived spectrum between species allowed us to eliminate the effect of the gradient (Figure 1) and rather focus on a potential effect of the life-history traits. As the simplest metric of the mutational spectrum, for each species we calculated first the Transition/Transversion ratio (Ts/Tv), as the sum of frequencies of all transitions divided by the sum of frequencies of all transversions. For 424 mammalian species with reconstructed Ts/Tv and known generation length, we observed a positive correlation between them (Supplementary Materials 2.3). Several additional analyses proved the robustness of this correlation: (i) we obtained the same trend with phylogenetic aware approach; (ii) we repeated the same trend splitting all species into several groups by quartiles of the generation length, by median of the generation length or by families; and finally (iii) we confirmed robustness of the results to the total number of polymorphisms used to calculate the mutational spectrum in different species (Supplementary Materials 2.3).

To understand further which substitution type(s) predominantly shaped the observed correlation between Ts/Tv and the generation length we performed twelve pairwise rank correlation analyses between each type of substitution and generation length. We observed that only A_H_>G_H_ frequency positively correlated with the generation length (Spearman’s rho = 0.252, nominal p-value = 1.188e-07) (Figure 2B), while several rare transversions showed a weak and negative correlation (T_H_>A_H_, T_H_>G_H_, C_H_>A_H_ and G_H_>T_H_: all Spearman’s rhos < -0.17, all nominal p-values are < 0.0003). The inclusion of all five types of these substitutions into a multiple linear model showed A_H_>G_H_ being the strongest component associated with generation length (Supplementary Materials 2.4). The observed effect was robust to phylogenetic inertia and the number of polymorphisms analyzed in each species (Supplementary Materials 2.4).

In order to analyze the mtDNA mutational spectra of mammals in an unsupervised way, we performed a dimensionality reduction by reconstructing principal components of mutational spectra of the 424 mammalian species. We observed that the first component is mainly driven by the most common C_H_>T_H_ substitutions, whereas the second is driven mainly by A_H_>G_H_ substitutions (Figure 2C left panel). We assessed if the first ten principal components were correlated with the generation length and observed that only the second one was significantly correlated with it (Figure 2C right panel, Supplementary Materials 2.5). Interestingly, the correlation of the second principal component with generation length was higher as compared to the sole effect of A_H_>G_H_ frequency, suggesting that the second component could reflect a complex nature of a longevity-associated mutational signature (Supplementary Materials 2.5). Additional analyses which take into account the effects of the total number of substitutions (Supplementary Materials 2.5), potential sequencing errors (Supplementary Materials 2.6) and nucleotide content (Supplementary Materials 2.7) confirmed the robustness of our results. Altogether, our analyses of within-species four-fold degenerate synonymous polymorphisms in hundreds of mammalian species demonstrated a robust association of mtDNA mutational spectrum (mainly the frequency of A_H_>G_H_) with the species-specific generation length.

### (3) mtDNA of mammals with high generation length are more A_H_ poor and G_H_ rich due to intensive A_H_>G_H_ mutagenesis

Mutational bias, if stronger than selection, in the long-term perspective is expected to change the genome-wide nucleotide content. Below we test this assumption.

First, in order to model the possible effect of the mutation bias on the nucleotide composition we used a computational simulation which, based on an input 12-component mutational spectrum, derived the expected neutral nucleotide composition. We run this simulation separately for mammals with very short-(less than the lower decile: 554 days, N = 27) and very long-(higher than the upper decile: 5221 days, N =25) generation length (Supplementary Materials 3.1); an average mutational spectrum of mammals with very long generation length was characterized by almost two-fold increased frequency of A_H_>G_H_ (Supplementary Materials 3.1). The results of these simulations demonstrated that an expected nucleotide composition of mammals with high generation length is characterized by the increased frequency of Gh and the decreased frequency of A_H_ (Supplementary Materials 3.1). In order to estimate how close the mammals are to their compositional nucleotide equilibrium we compared the expected nucleotide composition with the observed ones, which was derived using synonymous four-fold degenerate nucleotide content of twelve (all except ND6) protein-coding genes from the same species with very short and very long generations length (Figure 3A). We found that the observed nucleotide composition is rather similar with the expected one, which means that analyzed species are close enough to the compositional equilibrium. Moreover, we observed that species with short generation length tend to be closer to the expected equilibrium (see the horizontal dotted lines on Figure 3A) as compared to species with high generation length, probably because species with short generation length have an increased mutational rate (increased number of mtDNA replications per unit of time) and thus approach an equilibrium faster. Altogether, we observed that the synonymous four fold degenerate nucleotide composition of mammals is close to their neutral equilibrium and thus we expect to observe an effect of the mutation bias on the nucleotide content of mammalian mtDNA.

**FIGURE 3:**
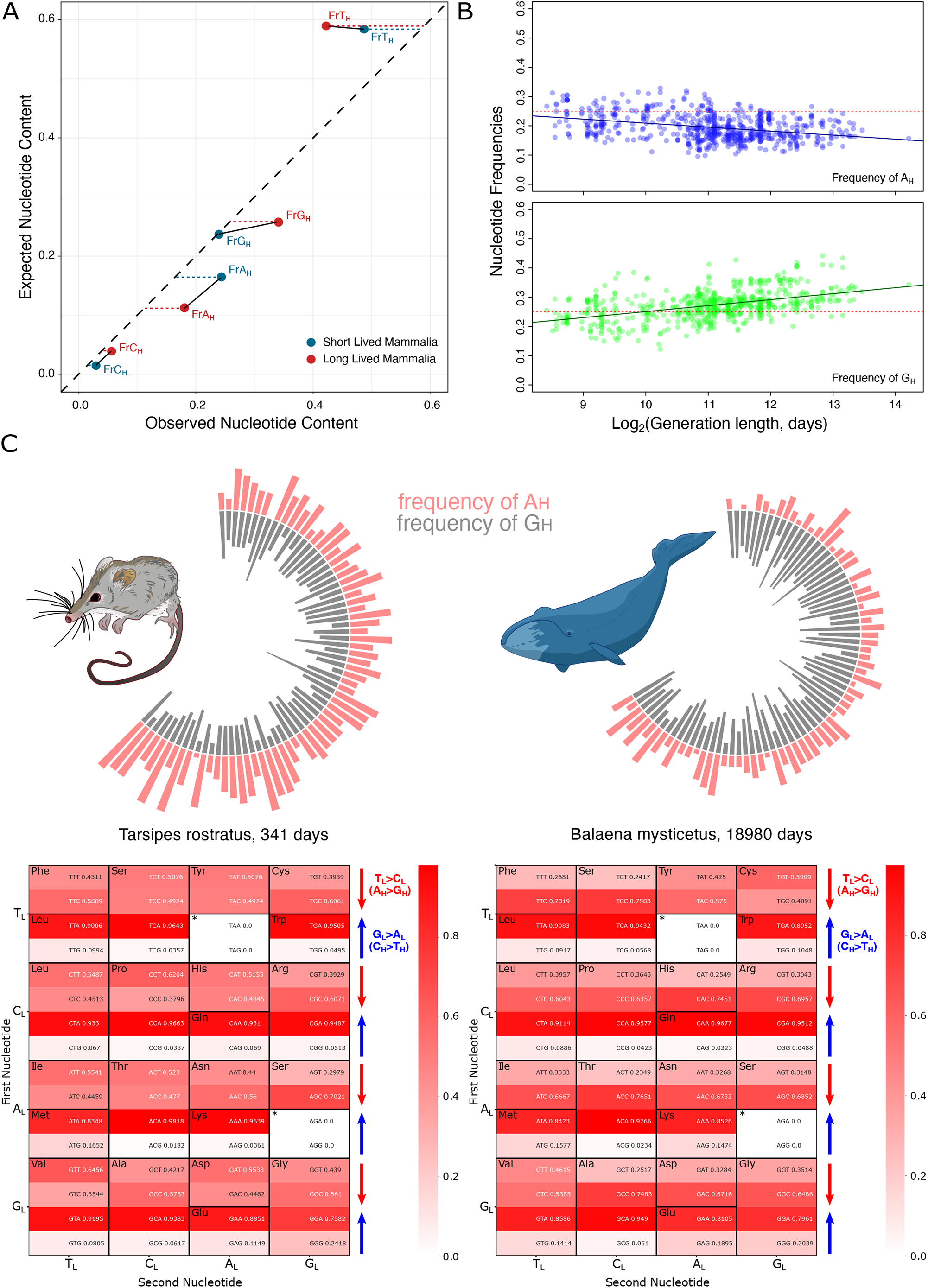
the long-term effect of the mutational bias: neutral nucleotide content in mammalian species 3A: A correlation between the expected (obtained in simulations, see supplementary materials 3.1) and observed neutral nucleotide content of mammals with very short and very long generation length. Due to an excess of A_H_>G_H_ substitutions in long-lived mammals (marked by red circles) they are more A_H_ poor and G_H_ rich for both expected and observed values. A location of all data points (red and blue circles) near the diagonal shows that mtDNA of mammals is close enough to a neutral equilibrium. However, short-lived species (marked by blue circles) are even closer to the diagonal (horizontal dotted blue line towards the diagonal are shorter than the red dotted lines), suggesting that they are evolving faster towards the neutral equilibrium. 3B. Nucleotide frequencies in neutral sites of all 13 protein-coding genes as a function of generation length - fraction of A_H_ is decreasing while fraction of G_H_ is increasing (N = 650). 3C. Structure of mtDNA of two mammalian species with extreme generation lengths: an opossum and a whale. Upper panel. Frequencies of A_H_ (red) and G_H_ (grey) nucleotides along the major arc of mtDNA of the most short-lived (opossum) and the most long-lived (whale) mammalian species from our dataset. Each bar represents the nucleotide frequency in a 20 nucleotide window. In both mammals A_H_ is decreasing and G_H_ is increasing along the major arc of mtDNA: from the bottom left (origin of replication of light strand) to the top right (origin of replication of heavy strand). However, additionally to the gradient, mtDNA of a whale has an integral, genome-wide, deficit of A_H_ and excess of G_H_ - a signature of an increased generation length. Lower panel. Heatmaps visualise asymmetry of the codon usage of 12 protein-coding genes (all except ND6). Whale is more contrast than opossum in terms of an asymmetry driven by the age-related T_L_>C_L_ (A_H_>G_H_) substitutions. Heatmaps of both species are equally contrasted in terms of an asymmetry driven by G_L_>A_L_ (C_H_>T_H_) substitutions, which have high and similar (not age related) substitution rate in both species.

Second, we tested if the increased A_H_>G_H_ (Fig 2) in species with high generation length would decrease frequencies of A_H_ and increase frequencies of G_H_ in the corresponding reference sequences? Since generation length correlates with the strength of the A_H_>G_H_ (Fig 2) we expect that generation length should demonstrate positive correlation with G_H_ and negative - with A_H_. Testing all four pairwise correlations between the species-specific generation length and the nucleotide content (A_H,_ T_H,_ G_H,_ C_H_) we observed two strongest correlations: negative with A_H_ and positive with G_H_ (Figure 3B; Supplementary Materials 3.2). Inclusion of all four types of nucleotide frequencies into the multiple linear model confirmed the importance of A_H_ and G_H_ only, the effect of which was also robust to the phylogenetic inertia (Supplementary Materials 3.2). Thus we concluded that mtDNA of mammals with long-versus short-generation length is more A_H_ poor and G_H_ rich (Figure 3B), which is in line with the more intensive A_H_>G_H_ mutagenesis in the former (Figure 2).

Third, we tested if an excess of G_H_ and deficit of A_H_ in long-lived species determines the positive G_H_A_H_ nucleotide skew. The G_H_A_H_ nucleotide skew approximates the level of asymmetry in the distribution of these two nucleotides and is calculated as (G_H_-A_H_)/(G_H_+A_H_). Based on four-fold degenerate synonymous positions of 12 genes (all except ND6) we estimated G_H_A_H_ skew for each mammalian species and correlated it with the generation length. As expected we obtained a positive correlation (phylogenetic generalized least squares: coefficient = 0.13, p-value = 2.9*10^−4^; see also Figure 3C). To visualize a contrast in G_H_A_H_ skew between the shortest- and the longest-lived species in our dataset we plotted A_H_ and G_H_ fractions along the major arc of mtDNA for opossum (generation length 341 days) and whale (generation length 18980 days) (Figure 3C). It is evident that on average opossum mtDNA has an excess of A_H_ (red color in Figure 3C) while whale has an excess of G_H_ (gray color in Figure 3C).

Fourth, we analyzed how A_H_>G_H_ mutagenesis affects asymmetry in codon usage. If A_H_>G_H_ is a strong and uniform mutational force, we expect that the majority of amino acids would demonstrate a deficit of XXA_H_ and excess of XXG_H_ codons. Because the light chain of mtDNA is equivalent to mRNA of 12 protein-coding genes (all except ND6) we analyze codon usage in terms on the light chain notation and thus, A_H_>G_H_ substitutions are expected to decrease frequency of XXT_L_ codons (because T_L_ is complementary to A_H_) and increase frequency of XXC_L_ codons (because C_L_ is complementary to G_H_). Similarly, C_H_>T_H_ mutations are expected to increase frequencies of XXA_L_ and decrease frequencies of XXG_L_ codons. To visualize this effect we plotted heatmaps of the codon usage of opossum and whale (based on 12 genes, all except ND6) (Figure 3C bottom panels). We observed that XXC_L_ asymmetry (see methods) is indeed stronger in a whale (0.70) as compared to an opossum (0.52). Interestingly, XXA_L_ asymmetry (see methods) is similarly high in both these species (whale: 0.93, opossum: 0.95). Quantifying the XXC_L_ asymmetry and XXA_L_ asymmetry across all mammalian species with complete mitochondrial genomes, we observed that (i) both XXC_L_ and XXA_L_ are significantly higher than the null expectation 0.5 (Supplementary Materials 3.3); (ii) XXA_L_ asymmetry is much stronger as compared to XXC_L_ asymmetry (Supplementary Materials 3.3), supporting that the C_H_>T_H_ substitution rate is much higher as compared to A_H_>G_H_ (see also Figure 2A) and (iii) XXC_L_ asymmetry correlates strongly and positively with the generation length of mammals, while XXA_L_ demonstrates weak negative correlation, which is not supported by the phylogeny aware statistics (Supplementary Materials 3.3), supporting that only A_H_>G_H_ substitutions (those which affect XXC_L_ asymmetry) are associated with the generation length.

Altogether we demonstrated, that A_H_>G_H_ mutagenesis, more pronounced in long-lived species, strongly shapes its reference sequences: nucleotide content (low A_H_ and high G_H_ frequency), nucleotide skew (srong positive G_H_A_H_ skew) and the codon usage (positive XXC_L_ asymmetry).

### (4) G_H_A_H_ nucleotide skew is a function of both time spent single stranded (TSSS) and the generation length

It has been shown that the frequency of A_H_>G_H_ substitutions depends on Time which parental heavy strand Spent in Single-Stranded (TSSS) condition during asynchronous mtDNA replication (Sanchez-Contreras et al. 2021; Faith and Pollock 2003). Genes, located close to the origin of light strand replication (O_L_), such as *COX1*, spend minimal time being single-stranded and demonstrate the low frequency of A_H_>G_H_, while genes located far away from O_L_ spend more time being single-stranded and demonstrate correspondingly higher frequencies of A_H_>G_H_ (Fig 1). Thus, we expect that effectively neutral nucleotide composition of mtDNA is a function of both: gene-specific TSSS and species-specific generation length. To test this we derived for each gene of each species the G_H_A_H_ skew and split all mammalian species into species with short and long generation length according to the median (median = 2190 days, N short = 325, N long = 319). Next we plotted G_H_A_H_ skew of mammals with short and long generation length for each gene, ranking them along the major arc from COX1 (rank equals 1) to CYTB (rank equals 10), corresponding to the increasing TSSS. As expected we observed that G_H_A_H_ skew increases with both gene-specific TSSS and species-specific generation length (Figure 4A). Performing multiple linear models, where G_H_A_H_ skew is a function of both TSSS and generation length, we confirmed that both factors affected the skew, moreover the effect sizes of TSSS and generation length were very similar (Supplementary Material 4.1).

**FIGURE 4:**
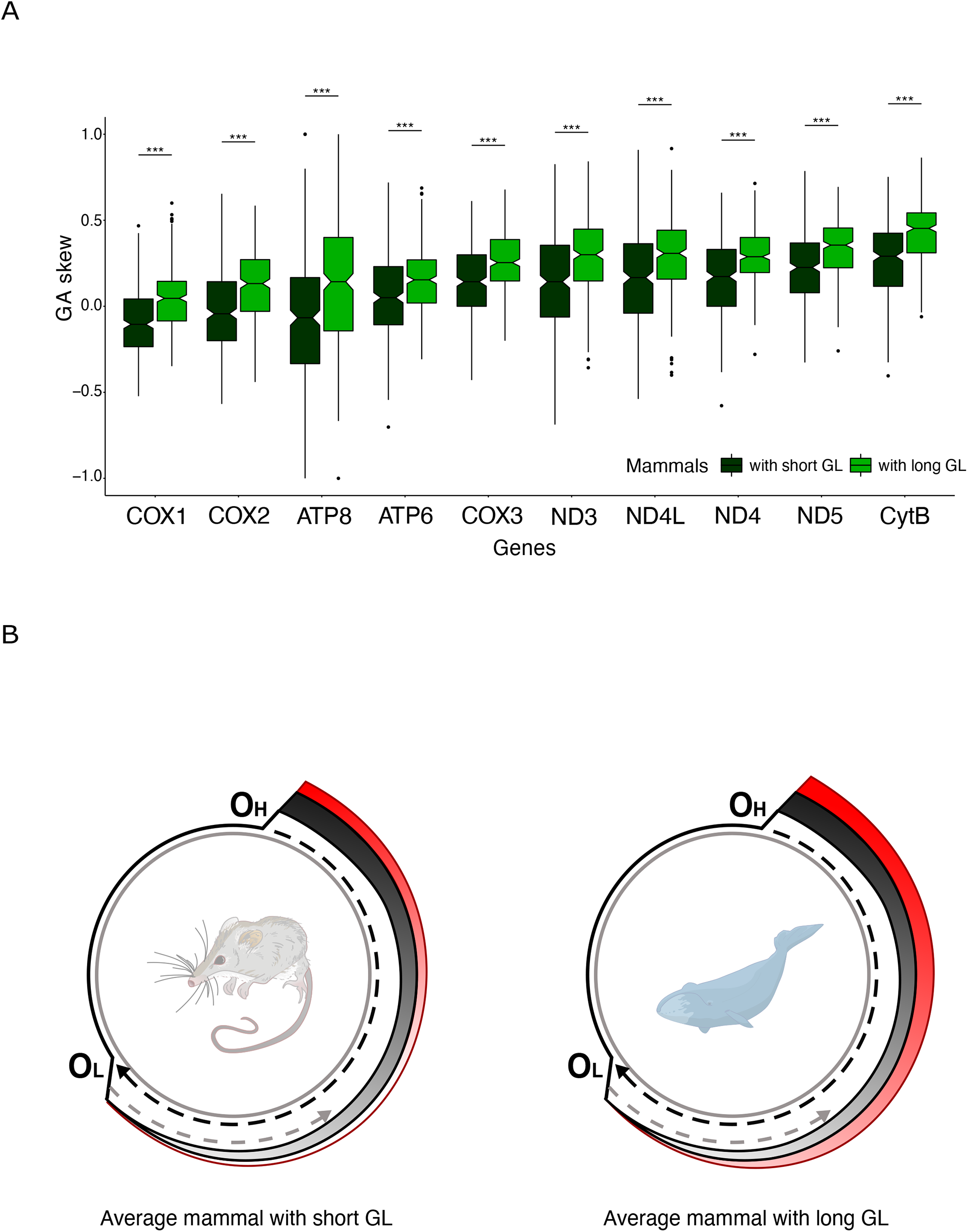
4A. Changes in nucleotide content along mtDNA of short- and long-lived mammals (N = 650). All genes (except for ND6) located in the major arc are ranked according to the time spent single stranded: from COX1 to CYTB. Pairs of boxplots for each gene represent G_H_A_H_ skew for short- and long-lived mammals splitted by the median generation length. G_H_A_H_ is increasing with both gene-specific TSSS and the species-specific generation length. (For statistical details see supplementary materials 4.1 and 4.2) 4B. A visual summary of the main finding: A _H_>G_H_ substitution rate (marked as red gradient) is increasing with both gene-specific TSSS and the species-specific generation length. The effect size of the GL is comparable with the effect size of TSSS (see supplementary materials 4.1). C_H_>T_H_ substitution rate (marked as grey gradient) is sensitive to TSSS only.

In the linear models we didn’t observe a significant interaction between TSSS and the generation length, suggesting that either these factors affect nucleotide composition independently of each other (Supplementary Materials 4.1) or interaction signal is too weak to be significant with our sample size. Our specific analyses, focused to uncover a potential interaction between TSSS and generation length indeed showed a positive trend, suggesting that mammals with high generation length demonstrate faster decrease in A_H_ and increase in G_H_ along the genome (Supplementary Materials 4.2, the same effect is visible on Figure 3B). Faster changes (stronger gradients) in A_H_ and G_H_ along the major arc of mtDNA in long-lived mammals can be interpreted as an interaction between TSSS and generation length as if substitution rate A_H_>G_H_ increases faster as a function of TSSS in case of high generation length.

Altogether, our results demonstrate that the nucleotide content, shaped by the mutational bias from A_H_ to G_H_, positively and strongly depends on both TSSS and the generation length.

## Discussion

We observed that A_H_>G_H_ substitution rate and its consequences such as nucleotide frequencies, G_H_A_H_ skew, codon usage asymmetry increase with organismal longevity. This finding was repeated on three rather different time scales: (i) months and years for somatic and de novo germline mutations (Fig 1); (ii) dozends or hundreds of thousands years for neutral within-species mtDNA polymorphisms (which is the average time of segregation of neutral within-species mtDNA polymorphisms (Atkinson, Gray, and Drummond 2008)) (Fig 2); (iii) millions of years for neutral mtDNA substitutions fixed between species (Fig 3-4). This universal trend requires a special explanation.

We consider that a process of mtDNA mutagenesis, rather than selection, is primarily responsible for all our findings. First, we expect no effect of selection in case of extremely rare, with variant allele frequency less than 1%, mtDNA mutations, called in duplex sequencing approach (Sanchez-Contreras et al. 2021) (Fig 1). Second, due to little or no evidence of selection on synonymous four-fold degenerate sites in mtDNA of mammals (Faith and Pollock 2003; Uddin and Chakraborty 2017) we consider the polymorphic variants (Fig 2) as well as the variants fixed between mammalian species (Fig 3-4) as effectively neutral.

We suggest that mtDNA mutagenesis is sensitive to the mitochondrial microenvironment which in turn depends on organism-specific generation length. Since oxidative damage is a main by-product of the aerobic metabolism and aging is associated with an increased oxidative damage, we propose that our key findings are driven by the oxidative-damage induced mutations.

It is important to emphasize, that our discovered substitution differs from the well-documented ROS-induced 7,8-Dihydro-8-oxo-20-deoxyguanosine (8-oxodG), leading to G > T transversions. It has been shown before that the expected hallmark of the oxidative damage, namely G>T substitutions, does not work in mtDNA: it is very rare, age-independent and demonstrates weak if any association with oxidative damage (Yuan et al. 2020; Kennedy et al. 2013; Hoekstra et al. 2016; Zsurka et al. 2018). Interestingly, G>T transversions (G_H_>T_H_ and C_H_>A_H_) in our work despite the rareness and weak effect, demonstrate an opposite to A_H_>G_H_ trend (Figure 2C left panel), which may reflect the classical oxidative damage signature (ROS-induced 8-oxodG) more pronounced in short-lived species due to their higher relative basal metabolic rate. However its effect indeed is weak.

Here, instead of G>T substitutions, we propose that the deamination of adenosine, which is the main source of A_H_>G_H_ substitutions, is associated with an oxidative damage of a single-stranded mtDNA. We propose that A>G is a novel mtDNA-marker of oxidative damage typical for a single-stranded DNA. Our results (Fig 1-4) and numerous papers, discussed below, support our hypothesis.

First, recent deep sequencing of de novo mtDNA mutations in aged versus young mice oocytes confirmed that an increased fraction of A_H_>G_H_ substitutions is the best mtDNA hallmark of oocyte aging (Arbeithuber et al. 2020). This discovery also suggests that continuous mtDNA turnover within the dormant oocytes can be a universal trait of all mammalian species (Arbeithuber et al. 2020) - and thus variation in mtDNA mutational spectra between different species can reflect the age-associated processes in different mammalian species.

Second, an excess of A:T>G:C substitutions has been observed recently in aerobically versus anaerobically grown Escherichia coli. It is important to emphasize that the signal was driven by the lagging strand, spending more time in single-stranded condition (Shewaramani et al. 2017). Results of the experiment are compatible with our hypothesis, that A>G transitions are associated with oxidative damage of single-stranded DNA.

Third line of evidence is coming from a mismatch repair (MMR) pathway, which can prevent mutations caused by oxidative stress (Bridge, Rashid, and Martin 2014). Mismatch repair-deficient mouse cell line shows a 6-50 fold increase in the rate of the A:T>G:C substitutions in the nuclear genome, and this increase is washed out in the presence of antioxidants (Shin and Turker 2002). Since MMR is absent in mtDNA, the level of oxidative damage in mtDNA can directly affects the rate of A>G transitions.

Fourth, it has been shown that A>G is the most asymmetric substitution in the human nuclear genome, which is a hallmark of a mutagen, inducing strand-specific DNA damage (Seplyarskiy et al. 2019). Despite the fact, that the key mutagen is still under question, we suggest that it can be associated with oxidative damage in case of mtDNA.

Fifth, it has been shown recently that the fraction of A_H_>G_H_ substitutions in mtDNA positively correlates with the ambient temperature of Actinopterygii species (Mikhailova et al. 2020). Because higher temperature is associated with an increased level of aerobic metabolism (Martin and Palumbi 1993) we assume that A_H_>G_H_ is a marker of oxidative damage.

Finally, A>G substitutions, associated with an oxidative damage, can be a key process, explaining a long-standing evolutionary puzzle of increased GC content of aerobic versus anaerobic bacteria (Naya et al. 2002; Romero et al. 2009; Aslam et al. 2019). According to the commonly accepted knowledge G>T is expected to be higher in aerobic versus anaerobic bacteria, making genomes of the formers more GC poor and AT rich. However numerous empirical evidence show the opposite: aerobic bacteria have increased GC content as compared to anaerobic (Naya et al. 2002; Romero et al. 2009; Aslam et al. 2019). Assuming that oxidative damage have similar mutagenic effects on both mitochondrial and bacterial genomes we hypothesize that extensive A>G substitution rate in aerobic bacteria can make their genomes more G rich.

An excess of G_H_ in mtDNA of long-lived mammals has been previously shown by Lehmann (Lehmann et al. 2006). He proposed a selective explanation of this observation assuming an increased stability of the G_H_-rich genomes which may confer an advantage to long-lived mammals. Our findings demonstrate that an excess of G_H_ in long-lived mammals may be a neutral consequences of A_H_>G_H_ mutagenesis, not selection (Fig 3-4). Additionally, low effective population size of long-lived mammals, increases the strength of the random genetic drift and the rate of fixation of slightly-deleterious variants in their mtDNA (K. Popadin et al. 2007; Nikolaev et al. 2007; K. Y. Popadin et al. 2013) making selective explanation even less probable. However, to evaluate the selective argument in more details we analysed the correlations of the nucleotide content (A, T, G, C) with generation length for each of three nucleotide positions in codons separately. We observed that A_H_ and G_H_ nucleotide frequencies at the third - the most neutral position of codons show the strongest correlations (negative for A_H_ and positive for G_H_) with generation length (supplementary materials 5), confirming that main factor, shaping an excess of G_H_ in long-lived mammals is mutagenesis, not selection.

The results of our study can significantly expand the usability of mtDNA mutational spectra. For example, low-heteroplasmy somatic mtDNA mutations from a merely neutral marker used to trace cellular lineages (Ludwig et al. 2019) can be transformed to a metric, associated with cell-specific aging state - a metric which can be especially important in highly heterogeneous tissues such as cancers. Low-heteroplasmy de novo germ-line mtDNA mutations (Arbeithuber et al. 2020) can predict the biological age of human oocytes, which can be used in *in vitro* fertilization techniques. MtDNA mutational spectra, reconstructed for non-model chordate species, may help to approximate their average generation length - a metric that is not always easier to estimate empirically.

Direct experimental investigation of the effect of different mutagens (Kucab et al. 2019) and especially an effect of oxidative damage (Degtyareva et al. 2019) and chemical modifications of nucleotides (C. W. Q. Koh et al. 2018, Hao et al. 2020) on single-stranded mtDNA will significantly improve our understanding of mtDNA mutagenesis.

Altogether, we demonstrated that A>G substitutions depend on both TSSS and GL and this relationship can be mediated by the sensitivity of this type of substitutions to the oxidative damage of the single-stranded DNA (Fig 4B).

## Methods

### Heavy chain notation of the 12-component mutational spectrum of mtDNA

Although it is traditional to refer to mtDNA substitutions with respect to the light strand, which corresponds to the reference mtDNA sequences, here we will refer to them based on the complementary heavy strand as several other authors (Raina et al. 2005; Krishnan et al. 2004). The heavy chain was the chain of choice since it is more mutagenic and thus the direction of the majority of the nucleotide substitutions would reflect the real mutagenic process (Yuan et al. 2020; Sanchez-Contreras et al. 2021; Kennedy et al. 2013). Hereafter, to simplify the biological interpretability of the mtDNA mutational spectrum, in the paper we use a 12-component spectrum based on heavy chain notation.

### Analysis of Duplex Sequencing mtDNA data

All data of somatic mtDNA mutations derifed from the duplex sequencing approach were obtained from the supplementary materials (Sanchez-Contreras et al. 2021). Control human mtDNA duplex sequencing data were used from two papers. In one paper age interval was 10-30 years (Baker et al. 2019), while on other the age interval was from 80 to 90 (Hoekstra et al. 2016).

### Reconstruction of the species-specific mutational spectrum for mammalian species

Using all available intraspecies sequences (at April 2016) of mitochondrial protein-coding genes we derived the mutational spectrum for each species. Briefly, we collected all available mtDNA sequences of any protein-coding genes for any chordate species, reconstructed the intraspecies phylogeny using an outgroup sequence (closest species for analyzed one), reconstructed ancestral states spectra in all positions at all inner tree nodes and finally got the list of single-nucleotide substitutions for each gene of each species. The pipeline is described in more details in Supplementary Materials 1.1. Using species with at least 15 single-nucleotide synonymous substitutions at four-fold degenerate sites we estimated mutational spectrum as a probability of each nucleotide to mutate into any other nucleotide (vector of 12 types of substitutions with sum equals one) for more than a thousand of chordata species. We normalized observed frequencies by nucleotide content in the third position of four-fold degenerative synonymous sites of a given gene. To eliminate the effect of nonuniform sampling of analyzed genes between different species the vast majority of our analyses were performed with *MT-CYB* gene - the most common gene in our dataset.

Generation length in days (as the average age of parents of the current cohort, reflecting the turnover rate of breeding individuals in a population) for mammals was downloaded from Pacifici et al. (Pacifici et al. 2013).

### Analyses of codon asymmetry

XXC_L_ codon asymmetry was defined as a median value of XXC_L_ /(XXC_L_ + XXT_L_) from each aminoacid. Similarly XXA_L_ asymmetry was defined as a median value of XXA_L_ /(XXA_L_ + XXG_L_).

### Analyses of complete mitochondrial genomes

We downloaded whole mitochondrial genomes from GenBank using the following search query: “Chordata”[Organism] AND (complete genome[All Fields] AND mitochondrion [All Fields] AND mitochondrion[filter]. We extracted non-overlapping regions of protein-coding genes, calculated codon usage and extracted fractions of A_H_, T_H_, G_H_ and C_H_ nucleotides in synonymous fourfold degenerate positions. For each protein-coding gene of each reference mitochondrial genome of mammalian species we estimated G_H_A_H_ skew as (G_H_-A_H_)/(G_H_+A_H_) using only synonymous fourfold degenerate sites.

### Data and scripts availability

Numerous methodological details are presented in supplementary materials 1-5. All statistical analyses were performed in R. All data and scripts are available in our public GitHub repository: https://github.com/polarsong/mtDNA_mutspectrum/.

## Acknowledgments

We thank Athanasios Kousathanas for statistical comments, the whole laboratory of Alexandre Reymond, Mikhail Gelfand and Vladimir Katanaev for discussions and valuable suggestions. K.P. was supported by the 5 Top 100 Russian Academic Excellence Project at the Immanuel Kant Baltic Federal University. E.O.T. is supported by a scholarship from the Austrian Science Fund (FWF, DOC 33-B27). This work was also supported by Russian Foundation of Basic Research [No. 19-29-04101 to I.M]. This work was also supported by Russian Science Foundation [No. 21-75-20143 to A.G., K.P, K.G. & V.Y.]

## Notes

### Competing Interest Statement

The authors have declared no competing interest.

## Literature

Alexandrov, Ludmil B., Serena Nik-Zainal, David C. Wedge, Samuel A J, Sam Behjati, Andrew V. Biankin, Graham R. Bignell, et al. 2013. “Signatures of Mutational Processes in Human Cancer.” Nature 500 (7463): 415–21.

Alexeyev, Mikhail F. 2009. “Is There More to Aging than Mitochondrial DNA and Reactive Oxygen Species?” FEBS Journal. https://doi.org/10.1111/j.1742-4658.2009.07269.x.

Arbeithuber, Barbara, James Hester, Marzia A. Cremona, Nicholas Stoler, Arslan Zaidi, Bonnie Higgins, Kate Anthony, Francesca Chiaromonte, Francisco J. Diaz, and Kateryna D. Makova. 2020. “Age-Related Accumulation of de Novo Mitochondrial Mutations in Mammalian Oocytes and Somatic Tissues.” PLoS Biology 18 (7): e3000745.

Aslam, Sidra, Xin-Ran Lan, Bo-Wen Zhang, Zheng-Lin Chen, Li Wang, and Deng-Ke Niu. 2019. “Aerobic Prokaryotes Do Not Have Higher GC Contents than Anaerobic Prokaryotes, but Obligate Aerobic Prokaryotes Have.” BMC Evolutionary Biology 19 (1): 35.

Atkinson, Q. D., R. D. Gray, and A. J. Drummond. 2008. “mtDNA Variation Predicts Population Size in Humans and Reveals a Major Southern Asian Chapter in Human Prehistory.” Molecular Biology and Evolution. https://doi.org/10.1093/molbev/msm277.

Baker, Kathryn T., Daniela Nachmanson, Shilpa Kumar, Mary J. Emond, Cigdem Ussakli, Teresa A. Brentnall, Scott R. Kennedy, and Rosa Ana Risques. 2019. “Mitochondrial DNA Mutations Are Associated with Ulcerative Colitis Preneoplasia but Tend to Be Negatively Selected in Cancer.” Molecular Cancer Research: MCR 17 (2): 488–98.

Belle, Elise M. S., Gwenael Piganeau, Mike Gardner, and Adam Eyre-Walker. 2005. “An Investigation of the Variation in the Transition Bias among Various Animal Mitochondrial DNA.” Gene 355 (August): 58–66.

Bridge, Gemma, Sukaina Rashid, and Sarah A. Martin. 2014. “DNA Mismatch Repair and Oxidative DNA Damage: Implications for Cancer Biology and Treatment.” Cancers 6 (3): 1597–1614.

Damuth, John. 1987. “Interspecific Allometry of Population Density in Mammals and Other Animals: The Independence of Body Mass and Population Energy-Use.” Biological Journal of the Linnean Society. https://doi.org/10.1111/j.1095-8312.1987.tb01990.x.

Degtyareva, Natalya P., Natalie Saini, Joan F. Sterling, Victoria C. Placentra, Leszek J. Klimczak, Dmitry A. Gordenin, and Paul W. Doetsch. 2019. “Mutational Signatures of Redox Stress in Yeast Single-Strand DNA and of Aging in Human Mitochondrial DNA Share a Common Feature.” PLOS Biology. https://doi.org/10.1371/journal.pbio.3000263.

Faith, Jeremiah J., and David D. Pollock. 2003. “Likelihood Analysis of Asymmetrical Mutation Bias Gradients in Vertebrate Mitochondrial Genomes.” Genetics 165 (2): 735–45.

Fraga, C. G., M. K. Shigenaga, J. W. Park, P. Degan, and B. N. Ames. 1990. “Oxidative Damage to DNA during Aging: 8-Hydroxy-2’-Deoxyguanosine in Rat Organ DNA and Urine.” Proceedings of the National Academy of Sciences of the United States of America 87 (12): 4533–37.

Guliaeva, N. A., E. A. Kuznetsova, and A. I. Gaziev. 2006. “[Proteins associated with mitochondrial DNA protect it against the action of X-rays and hydrogen peroxide].” Biofizika 51 (4): 692–97.

Hao, Ziyang, Tong Wu, Xiaolong Cui, Pingping Zhu, Caiping Tan, Xiaoyang Dou, Kai-Wen Hsu, et al. 2020. “N-Deoxyadenosine Methylation in Mammalian Mitochondrial DNA.” Molecular Cell 78 (3): 382–95.e8.

Harris, Kelley, and Jonathan K. Pritchard. 2017. “Rapid Evolution of the Human Mutation Spectrum.” eLife 6 (April). https://doi.org/10.7554/eLife.24284.

Hebert, Paul D. N., Alina Cywinska, Shelley L. Ball, and Jeremy R. deWaard. 2003. “Biological Identifications through DNA Barcodes.” Proceedings. Biological Sciences / The Royal Society 270 (1512): 313–21.

Hoekstra, Jake G., Michael J. Hipp, Thomas J. Montine, and Scott R. Kennedy. 2016. “Mitochondrial DNA Mutations Increase in Early Stage Alzheimer Disease and Are Inconsistent with Oxidative Damage.” Annals of Neurology 80 (2): 301–6.

Ju, Young Seok, Ludmil B. Alexandrov, Moritz Gerstung, Inigo Martincorena, Serena Nik-Zainal, Manasa Ramakrishna, Helen R. Davies, et al. 2014. “Origins and Functional Consequences of Somatic Mitochondrial DNA Mutations in Human Cancer.” eLife 3 (October). https://doi.org/10.7554/eLife.02935.

Kennedy, Scott R., Jesse J. Salk, Michael W. Schmitt, and Lawrence A. Loeb. 2013. “Ultra-Sensitive Sequencing Reveals an Age-Related Increase in Somatic Mitochondrial Mutations That Are Inconsistent with Oxidative Damage.” PLoS Genetics 9 (9): e1003794.

Koh, Casslynn W. Q., Yeek Teck Goh, Joel D. W. Toh, Suat Peng Neo, Sarah B. Ng, Jayantha Gunaratne, Yong-Gui Gao, Stephen R. Quake, William F. Burkholder, and Wee Siong S. Goh. 2018. “Single-Nucleotide-Resolution Sequencing of Human N6-Methyldeoxyadenosine Reveals Strand-Asymmetric Clusters Associated with SSBP1 on the Mitochondrial Genome.” Nucleic Acids Research 46 (22): 11659–70.

Koh, Gene, Andrea Degasperi, Xueqing Zou, Sophie Momen, and Serena Nik-Zainal. 2021. “Mutational Signatures: Emerging Concepts, Caveats and Clinical Applications.” Nature Reviews. Cancer 21 (10): 619–37.

Kucab, Jill E., Xueqing Zou, Sandro Morganella, Madeleine Joel, A. Scott Nanda, Eszter Nagy, Celine Gomez, et al. 2019. “A Compendium of Mutational Signatures of Environmental Agents.” Cell 177 (4): 821–36.e16.

Lehmann, Gilad, Arie Budovsky, Khachik K. Muradian, and Vadim E. Fraifeld. 2006. “Mitochondrial Genome Anatomy and Species-Specific Lifespan.” Rejuvenation Research 9 (2): 223–26.

Ludwig, Leif S., Caleb A. Lareau, Jacob C. Ulirsch, Elena Christian, Christoph Muus, Lauren H. Li, Karin Pelka, et al. 2019. “Lineage Tracing in Humans Enabled by Mitochondrial Mutations and Single-Cell Genomics.” Cell 176 (6): 1325–39.e22.

Martin, A. P., and S. R. Palumbi. 1993. “Body Size, Metabolic Rate, Generation Time, and the Molecular Clock.” Proceedings of the National Academy of Sciences of the United States of America 90 (9): 4087–91.

Mikhailova, Alina G., Victor Shamansky, Alina A. Mikhailova, Kristina Ushakova, Evgenii Tretyakov, Sergey Oreshkov, Dmitry Knorre, et al. 2020. “A Mitochondrial Mutational Signature of Temperature and Longevity in Ectothermic and Endothermic Vertebrates.” Cold Spring Harbor Laboratory. https://doi.org/10.1101/2020.07.25.221184.

Montooth, Kristi L., and David M. Rand. 2008. “The Spectrum of Mitochondrial Mutation Differs across Species.” PLoS Biology 6 (8): e213.

Moorjani, Priya, Carlos Eduardo G. Amorim, Peter F. Arndt, and Molly Przeworski. 2016. “Variation in the Molecular Clock of Primates.” https://doi.org/10.1101/036434.

Naya, Hugo, Héctor Romero, Alejandro Zavala, Beatriz Alvarez, and Héctor Musto. 2002. “Aerobiosis Increases the Genomic Guanine plus Cytosine Content (GC%) in Prokaryotes.” Journal of Molecular Evolution 55 (3): 260–64.

Nikolaev, Sergey I., Juan I. Montoya-Burgos, Konstantin Popadin, Leila Parand, Elliott H. Margulies, Stylianos E. Antonarakis, National Institutes of Health Intramural Sequencing Center Comparative Sequencing Program, and Others. 2007. “Life-History Traits Drive the Evolutionary Rates of Mammalian Coding and Noncoding Genomic Elements.” Proceedings of the National Academy of Sciences 104 (51): 20443–48.

Ollason, J. G. 1987. “R. H. Peters 1986. The Ecological Implications of Body Size. Cambridge University Press, Cambridge. 329 Pages. ISBN 0-521-2886-X. Price: £12.50, US$16.95 (paperback).” Journal of Tropical Ecology. https://doi.org/10.1017/s0266467400002224.

Pacifici, Michela, Luca Santini, Moreno Di Marco, Daniele Baisero, Lucilla Francucci, G. Grottolo Marasini, Piero Visconti, and Carlo Rondinini. 2013. “Generation Length for Mammals.” Nature Conservation 5: 87–94.

Popadin, Konstantin, Leonard V. Polishchuk, Leila Mamirova, Dmitry Knorre, and Konstantin Gunbin. 2007. “Accumulation of Slightly Deleterious Mutations in Mitochondrial Protein-Coding Genes of Large versus Small Mammals.” Proceedings of the National Academy of Sciences of the United States of America 104 (33): 13390–95.

Popadin, Konstantin Yu, Sergey I. Nikolaev, Thomas Junier, Maria Baranova, and Stylianos E. Antonarakis. 2013. “Purifying Selection in Mammalian Mitochondrial Protein-Coding Genes Is Highly Effective and Congruent with Evolution of Nuclear Genes.” Molecular Biology and Evolution 30 (2): 347–55.

Raina, Sameer Z., Jeremiah J. Faith, Todd R. Disotell, Hervé Seligmann, Caro-Beth Stewart, and David D. Pollock. 2005. “Evolution of Base-Substitution Gradients in Primate Mitochondrial Genomes.” Genome Research 15 (5): 665–73.

Reyes, A., C. Gissi, G. Pesole, and C. Saccone. 1998. “Asymmetrical Directional Mutation Pressure in the Mitochondrial Genome of Mammals.” Molecular Biology and Evolution. https://doi.org/10.1093/oxfordjournals.molbev.a026011.

Romero, Héctor, Emiliano Pereira, Hugo Naya, and Héctor Musto. 2009. “Oxygen and Guanine-Cytosine Profiles in Marine Environments.” Journal of Molecular Evolution 69 (2): 203–6.

Sanchez-Contreras, Monica, Mariya T. Sweetwyne, Brendan F. Kohrn, Kristine A. Tsantilas, Michael J. Hipp, Elizabeth K. Schmidt, Jeanne Fredrickson, et al. 2021. “A Replication-Linked Mutational Gradient Drives Somatic Mutation Accumulation and Influences Germline Polymorphisms and Genome Composition in Mitochondrial DNA.” Nucleic Acids Research 49 (19): 11103–18.

Sato, Ken, and Miyuki Sato. 2017. “Multiple Ways to Prevent Transmission of Paternal Mitochondrial DNA for Maternal Inheritance in Animals.” Journal of Biochemistry 162 (4): 247–53.

Seplyarskiy, Vladimir B., Evgeny E. Akkuratov, Natalia Akkuratova, Maria A. Andrianova, Sergey I. Nikolaev, Georgii A. Bazykin, Igor Adameyko, and Shamil R. Sunyaev. 2019. “Error-Prone Bypass of DNA Lesions during Lagging-Strand Replication Is a Common Source of Germline and Cancer Mutations.” Nature Genetics 51 (1): 36–41.

Shewaramani, Sonal, Thomas J. Finn, Sinead C. Leahy, Rees Kassen, Paul B. Rainey, and Christina D. Moon. 2017. “Anaerobically Grown Escherichia Coli Has an Enhanced Mutation Rate and Distinct Mutational Spectra.” PLoS Genetics 13 (1): e1006570.

Shin, Chi Y., and Mitchell S. Turker. 2002. “A:T → G:C Base Pair Substitutions Occur at a Higher Rate than Other Substitution Events in Pms2 Deficient Mouse Cells.” DNA Repair. https://doi.org/10.1016/s1568-7864(02)00149-0.

Tacutu, Robi, Thomas Craig, Arie Budovsky, Daniel Wuttke, Gilad Lehmann, Dmitri Taranukha, Joana Costa, Vadim E. Fraifeld, and João Pedro de Magalhães. 2013. “Human Ageing Genomic Resources: Integrated Databases and Tools for the Biology and Genetics of Ageing.” Nucleic Acids Research 41 (Database issue): D1027–33.

Tanaka, M., and T. Ozawa. 1994. “Strand Asymmetry in Human Mitochondrial DNA Mutations.” Genomics 22 (2): 327–35.

Uddin, Arif, and Supriyo Chakraborty. 2017. “Synonymous Codon Usage Pattern in Mitochondrial CYB Gene in Pisces, Aves, and Mammals.” Mitochondrial DNA. Part A, DNA Mapping, Sequencing, and Analysis 28 (2): 187–96.

Von Stetina, Jessica R., and Terry L. Orr-Weaver. 2011. “Developmental Control of Oocyte Maturation and Egg Activation in Metazoan Models.” Cold Spring Harbor Perspectives in Biology 3 (10): a005553.

Yuan, Yuan Yuan, PCAWG Consortium, Young Seok Ju, Youngwook Kim, Jun Li, Yumeng Wang, et al. 2020. “Comprehensive Molecular Characterization of Mitochondrial Genomes in Human Cancers.” Nature Genetics. https://doi.org/10.1038/s41588-019-0557-x.

Zou, Xueqing, Gene Ching Chiek Koh, Arjun Scott Nanda, Andrea Degasperi, Katie Urgo, Theodoros I. Roumeliotis, Chukwuma A. Agu, et al. 2021. “A Systematic CRISPR Screen Defines Mutational Mechanisms Underpinning Signatures Caused by Replication Errors and Endogenous DNA Damage.” Nature Cancer 2 (6): 643–57.

Zsurka, Gábor, Viktoriya Peeva, Alexander Kotlyar, and Wolfram S. Kunz. 2018. “Is There Still Any Role for Oxidative Stress in Mitochondrial DNA-Dependent Aging?” Genes 9 (4). https://doi.org/10.3390/genes9040175.

