## Supplementary Materials for "A mitochondria-specific mutational signature of aging: increased rate of A>G substitutions on a heavy chain"

Supplementary Materials to the manuscript of Mikhailova et al. "A mitochondria-specific mutational signature of aging: increased rate of A>G substitutions on heavy chain"

***(1) Frequency of de novo $A_H$>$G_H$ mutations increases with age in soma and germ-line***

(1.1) Ah>Gh is the most age-sensitive transition in mtDNA

*github:*
[https://github.com/polarsong/mtDNA_mutspectrum/blob/DuplexSeqData/Head/2Scripts/DuplexSeqDataAnalyses01.R](https://github.com/polarsong/mtDNA_mutspectrum/blob/DuplexSeqData/Head/2Scripts/DuplexSeqDataAnalyses01.R)

*Table S1. Results of the linear models, describing frequencies of the two most common mutations as a function of bins of the time spent single stranded (TSSS).* Ah>Gh is the most age-sensitive transition: an increase in the slopes from young mice to old mice to humans is the strongest (from 0.021 to 0.127 to 1.482).

| **age category** | **mutation type** | **intercept** | **slope** | **p-value** |
|---|---|---|---|---|
| young mice | Ch>Th | 20.273*** | 0.355 | <0.001*** |
| young mice | Ah>Gh | 1.736*** | 0.021 | <0.001*** |
| old mice | Ch>Th | 83.267*** | 0.703 | <0.001*** |
| old mice | Ah>Gh | 3.915*** | 0.127 | <0.001*** |
| humans | Ch>Th | 331.135*** | 4.866 | <0.001*** |
| humans | Ah>Gh | 100.536*** | 1.482 | <0.001*** |

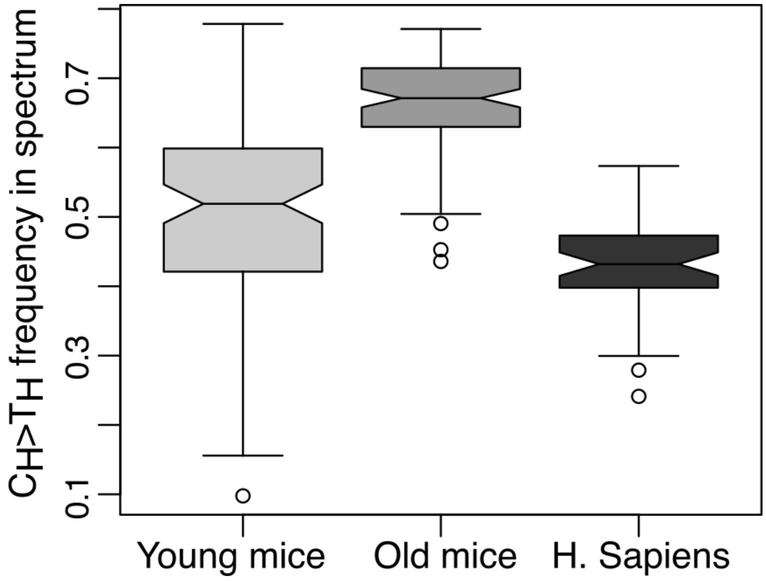


*Figure S1. The fraction of Ch>Th in the total mtDNA mutational spectrum shows no trends (increasing or decreasing) with age. P-values from all three pairwise comparisons are less than 1.583e-05, Mann-Whitney U test. Only Ah>Gh demonstrated the increasing trend (Fig 1C, lower panel).*

(1.2) mtDNA mutational spectrum is changing with female reproductive age: Ah>Gh is the best marker of oocyte age

In order to test potential changes in mtDNA mutational spectra during oocyte aging, we analyzed data from two papers dedicated to de novo mtDNA mutations in humans. Having access to the information about mothers' age at the moment of giving birth from studies of Rebolledo-Jaramillo (Rebolledo-Jaramillo et al. 2014) we observed that $A_H$>$G_H$ de novo mutations were characterized by the highest age of reproduction. Having access to the variant allele frequency (VAF) values of de novo mtDNA mutations from the study of Wei et al 2020 (Wei et al. 2019) we observed the later onset of $A_H$>$G_H$ as compared to other mutations. These results are in line with our previous findings. Details of the analyses are below.

Analysis of data from Rebolledo et al (Rebolledo-Jaramillo et al. 2014). Ah>Gh is characterized by the highest age of fertilization (average age of fertilization is 34.1, N = 4) as compared to Ch>Th (average age of fertilization is 29.9, N = 4) or all other substitutions (average age of fertilization is = 32.1, N = 12). Due to the small sample size (16 mutations in total) this trend was not significant (p = 0.19 if we compare Ah>Gh with all other substitutions and p = 0.0956 if we compare Ah>Gh with Ch>Th; one-sided Mann-Whitney U test).

*github*:
https://github.com/polarsong/mtDNA_mutspectrum/blob/Humans/Head/2Scripts/Humans.RebolledoAnalyses.R

Analysis of data from Wei et al (Wei et al. 2019). Data on the age of reproduction were not available and we used VAF of each de novo mtDNA variant as a function of oocyte age. We expect that AH>GH mutations occur mainly in aged oocytes and thus are characterized by lower VAF as compared to other substitutions. To decrease potential selection effects we concentrated only on rare (with VAF < 10%) substitutions. Next, subsetting offspring with at least two rare de novo mtDNA transitions, one being AH>GH and another non AH>GH, (23 offspring in total) we observed that AH>GH on average has 1% lower VAF as compared to non AH>GH. The statistical difference between VAF(AH>GH) and VAF(non AH>GH) was marginally significant (p = 0.05015, paired one-sided Mann-Whitney U test).

*github*:
https://github.com/polarsong/mtDNA_mutspectrum/blob/Humans/Head/2Scripts/Wei2019OffspringMutTypesComparison.R

**(2) Ah>Ghs are more prevalent in mammals with high generation length: an evidence from polymorphism-derived mutational spectra**

(2.1) The pipeline to derive mtDNA mutational spectra based on intraspecies polymorphisms

The most important pipeline steps were the following:
(i) The generation of the local nucleotide BLAST-database based on all available nucleotide sequences of mitochondrial protein-coding genes;
(ii) An retrieving of the intraspecies sequences from this database using *tblastn* software (ncbi-blast 2.6.0+ package) and RefSeq protein query sequence of a given species and a given mitochondrial gene;
(iii) An alignment of all mtDNA sequences for a given species and a given gene using *macse* v1.01b. This software performs codon-based multiple alignment (we used standard mitochondrial genetic code), and allows to check for inner stop codons in analyzed sequences (we selected out such positions from alignment);
(iv) Reconstruction of the tree. Unrooted tree topology of intraspecies sequences was derived by *RaxML* v.8.2.9 and "-m GTRGAMMAIX" option, we used best tree from 50 alternative runs on distinct starting trees ("-N 50" option) for selecting best tree topology. After that tree was rooted by nearest neighbor sequence from other species. This nearest neighbor sequence was found as a first blast hit followed by sequences of given species in our sequence database;
(v) Reconstruction of the ancestral sequences in each inner tree node. The reconstruction of intraspecies ancestral sequences was based on maximum likelihood tree topology and was made using two alternative approaches: maximum parsimony and maximum likelihood. Maximum parsimony ancestral reconstruction was done using *dnapars* program of *phylip* v.3.697 package. Maximum likelihood reconstruction (marginal ancestral states reconstruction) was done using *RaxML* v.8.2.9 and "-m GTRGAMMAIX -f A" options. For ancestral reconstruction we used nucleotide-based (not codon based) substitution models because in our downstream analyses we used only four-fold degenerate sites;
(vi) The list of polarized single-nucleotide substitutions occurred within the four-fold degenerate sites were used to reconstruct the mutational spectrum in R. See below also the Normalization step (Supplementary Materials 2.2)

(2.2) Normalisation of species-specific mtDNA mutational spectrum

*github:*
*https://github.com/polarsong/mtDNA_mutspectrum/blob/VertebratePolymorphisms/Body/3Results/VertebratePolymorphisms.MutSpecData.OnlyFourFoldDegAllGenes.txt - note that all metadata is encoded in ligth chain notation*

The fractions of observed substitutions depend on the frequencies of ancestral nucleotides in the synonymous fourfold degenerate sites, which are highly non-uniform. After normalizing the observed nucleotide substitutions (Figure S2A left panel) by the ancestral nucleotide frequency (Figure S2A right panel) we obtained a mutational spectrum as a probability of a given nucleotide to mutate to any other nucleotide irrespective of its frequency. Each species-specific mutational spectrum, presented as a vector of twelve substitution rates was additionally transformed to frequencies so that the total sum of all the substitution rates equals to one.

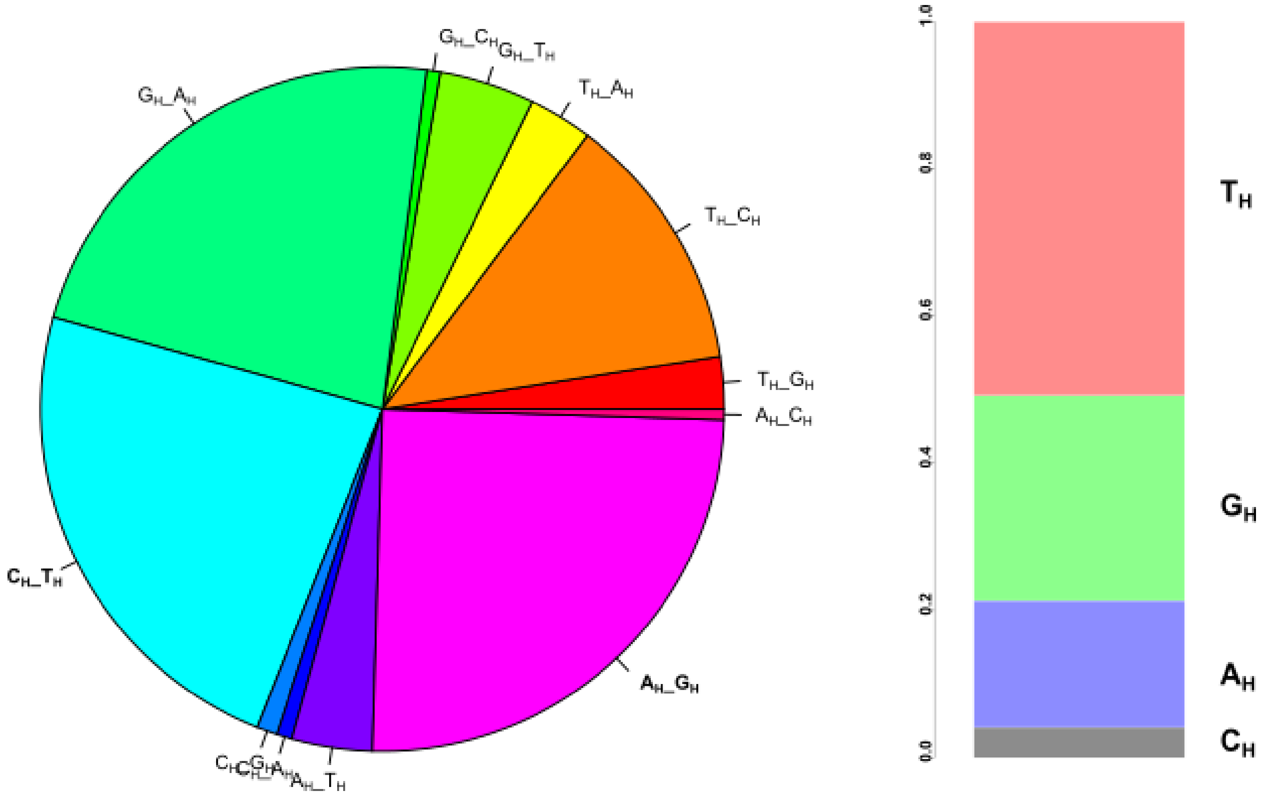


*Figure S2A. Derivation of mtDNA mutational spectrum for mammalian species (N = 611).*
*Left panel: observed frequencies of twelve types of nucleotide substitutions in four fold degenerate synonymous sites of all available mtDNA protein-coding genes.*
*Right panel: nucleotide content in four fold degenerate synonymous sites of all available mtDNA protein-coding genes.*
*After the normalization of the 12 substitution rates, depicted on the left panel, on the nucleotide content, depicted on the right panel, we obtained the normalized spectrum (Fig 1A) used in downstream analyses. For example, substitution rate of Th>Ch significantly decreased after normalization on high nucleotide content Th, while the fraction of Ch>Th increased after the normalization on very small number of Ch.*

(2.3) Species-specific Ts/Tv increases with the generation length

*Github with the generation length database:*
[*https://github.com/polarsong/mtDNA_mutspectrum/blob/VertebratePolymorphisms/Body/1Raw/GenerationLenghtforMammals.xlsx.txt*](https://github.com/polarsong/mtDNA_mutspectrum/blob/VertebratePolymorphisms/Body/1Raw/GenerationLenghtforMammals.xlsx.txt)

We observed a positive correlation between Ts/Tv and the species-specific generation length (Spearman's rho = 0.23, p-value = 2.021e-06, N = 424).

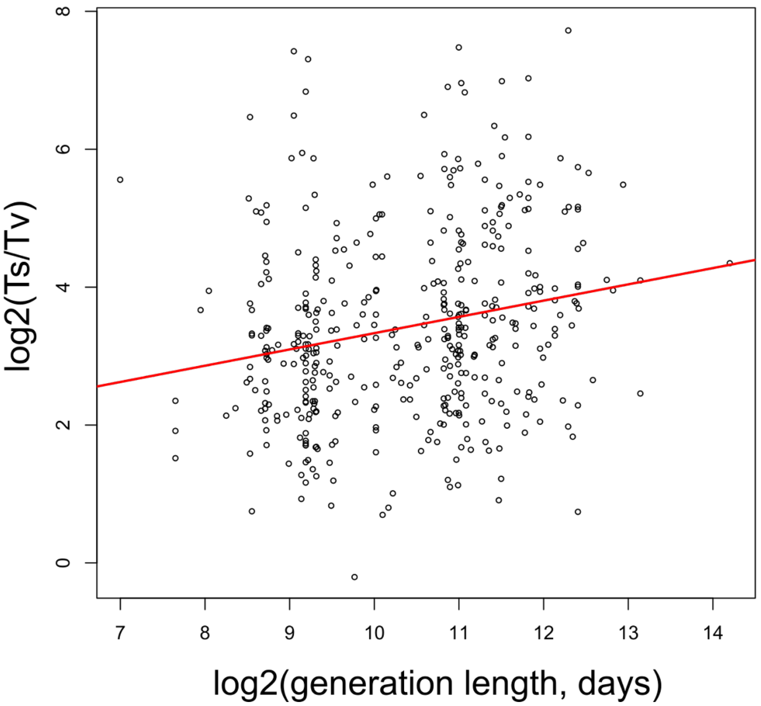


*Figure S2B. Positive correlation between generation length and Ts/Tv in mammals*

To prove robustness of our results we confirmed that an increase in Ts/Tv with the generation length is being observed also when we split all mammals into several groups:
(i) by quartiles of the generation length (Fig S2C);
(ii) by median of the generation length (Fig S2D) and
(iii) by families (Table S2A, Fig S2E).

Quartile split. When we split mammalian species into four quartiles, Ts/Tv is increasing from lower to higher quartiles: all possible pairwise comparisons of quartiles, except the comparison of the first and the second ones, show significant difference in Ts/Tv (Fig S2C, Mann-Whitney U test, all p-values < 0.05).

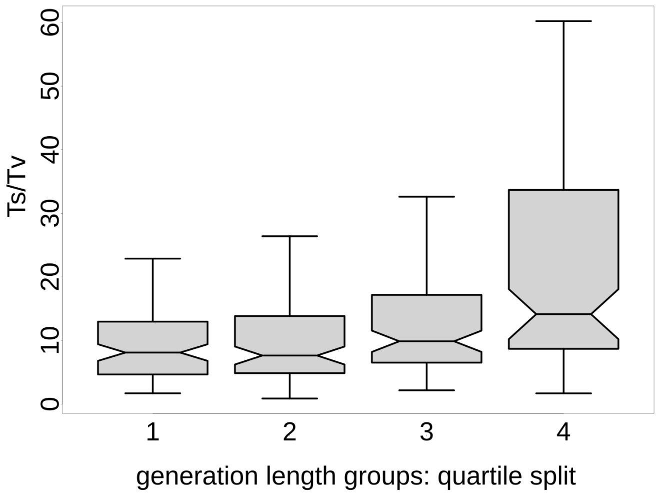


*Figure S2C. Mammalian species with high generation length demonstrate increased Ts/Tv (groups are split by quartiles)*

Median split. The median split of mammalian species by the generation length (median = 1497 days) also shows higher Ts/Tv in long- versus short-lived mammals (Fig S2D, p-value = 1.687e-06, Mann-Whitney U test).

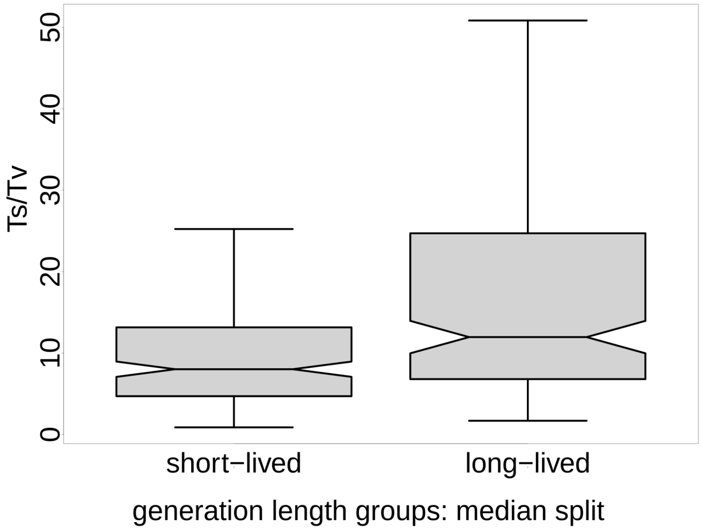


*Figure S2D. Mammalian species with high generation length demonstrate increased Ts/Tv (groups are split by a median)*

Family split. An increase in Ts/Tv with generation length is also pronounced on a family level (Table S2A). For each mammalian family, containing at least 3 species in our dataset, we estimated median Ts/Tv and median of generation length. Using Spearman rank correlation we confirmed a positive trend between these values (p = 0.0016, Spearman Rho = 0.85, N = 11). In the table below (Table S2A) there are descriptive statistics of eleven families and double boxplots (*Figure S2E*) visualize the distribution of Ts/Tv and generation length in each family.

*Table S2A. Family-based descriptive statistics of the analyzed dataset: median Ts/Tv and median of generation length for each family. The abbreviations introduced in the table are  used in the figure S2E below.*

| Family | median Ts/Tv | median generation length (in days) | abbreviation | number of species |
|---|---|---|---|---|
| Insectivora | 8.81 | 427 | Ins | 41 |
| Rodentia | 7.46 | 601 | Rod | 120 |
| Didelphimorphia | 7.08 | 643 | Did | 18 |
| Lagomorpha | 6.55 | 1047 | Lag | 19 |
| Chiroptera | 10.75 | 2064 | Chi | 81 |
| Carnivora | 13.21 | 2435 | Car | 34 |
| Cervidae | 13.33 | 2555 | Cer | 7 |
| Suidae | 10.57 | 2606 | Sui | 4 |
| Bovidae | 13.34 | 2870 | Bov | 16 |
| Primates | 14.14 | 3806 | Pri | 49 |
| Cetacea | 27.74 | 5158 | Cet | 6 |

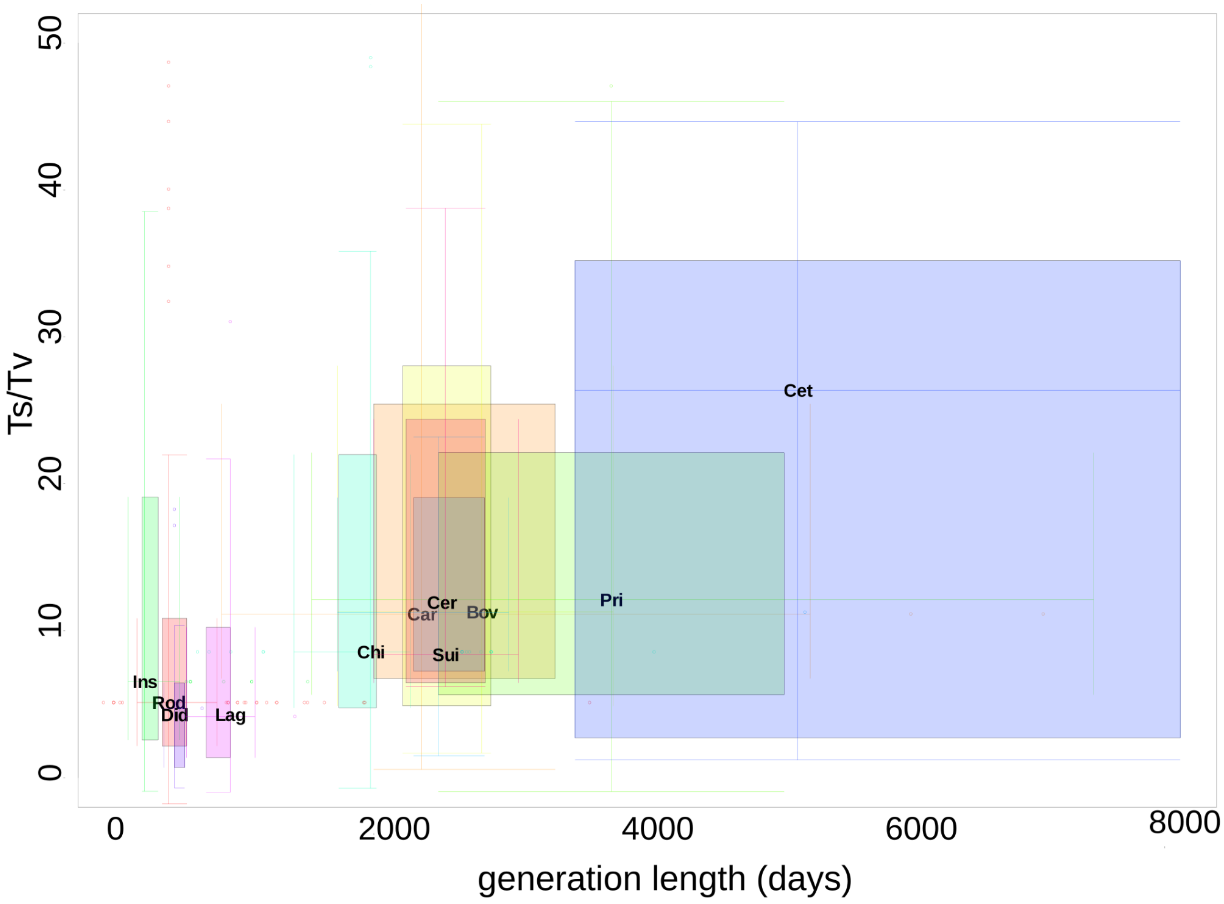


*Figure S2E. double boxplots visualize the distribution of Ts/Tv and generation length in each family. Abbreviations are as in Table S2A.*

Boxplots were obtained using the R library "boxplotdbl". The filled surface for each family reflects interquartile range, cross of lines represents medians, whiskers mark minimum and maximum, excluding outliers (some outliers are not shown on the plot). A position of the three-letter abbreviation for each family corresponds to medians of the generation length and Ts/Tv of the corresponding family.

<u>Phylogenetic Generalised Least-Squares regression (PGLS).</u> Although several analyses above demonstrated the robustness of the association between Ts/Tv and generation length we also performed phylogenetic generalised least-squares regression (PGLS) implemented in R library "caper". Due to the necessity to merge our dataset with an available phylogenetic tree we kept less than a half of our species (N = 211), but despite this fact the main trend is still significant. First, we demonstrated that Ts/Tv positively correlates with generation length (model 2.3A in the table S2B). Second, we added to the model the number of mutations as a number of reconstructed synonymous four-fold degenerate substitutions in CYB, which were used to reconstruct species-specific mutational spectrum (model 2.3B in the table S2B). Despite the fact that the number of mutations negatively correlates with Ts/Tv (probably due to higher sample size of short-lived mammals), the effect of the generation length on Ts/Tv stayed significant. Third, noting the intercept non-significantly deviates from the zero in model 2.3B we rerun the same model through the origin (fixing intercept at zero) and obtained strong and significant correlation between Ts/Tv and generation length (model 2.3C in the table S2B). In the final model 2.3C generation length is associated with Ts/Tv much more significantly as compared to the 'number of mutations'.

*Table S2B. Phylogenetic generalised least-squares regression (PGLS) models, describing the relationships between Ts/Tv and generation length*

| ***model*** | ***variable*** | ***coefficients*** | ***p values*** |
|---|---|---|---|
| ***2.3A: Ts/Tv ~ log2(generation length)****;* *N=211, lambda [ ML] = 0.000* | *intercept* | *-26.4047* | *0.094182* |
| | *log2(generation length)* | *4.1771* | *0.004472* |
| ***2.3B: Ts/Tv ~ log2(generation length) + log2(number of mutations)****;* *N=211, lambda [ ML] = 0.000* | *intercept* | *12.2181* | *0.558976* |
| | *log2(generation length)* | *2.9679* | *0.048715* |
| | *log2(number of mutations)* | *-4.3222* | *0.006407* |
| ***2.3C: Ts/Tv ~ 0 + log2(generation length) + log2(number of mutations)****;* *N=211, lambda [ ML] = 0.000* | *log2(generation length)* | *3.75742* | *2.454e-08* |
| | *log2(number of mutations)* | *-3.70511* | *0.001629* |

(2.4) A fraction of $A_H$>$G_H$ increases in species with high generation length

*github:*
https://github.com/polarsong/mtDNA_mutspectrum/tree/VertebratePolymorphisms/Head/2Scripts/*VertebratePolymorphisms.MutSpecComparisons.Analyses.Ecology.Mammals.R*

We included all five types of substitutions ($A_H$>$G_H$, $T_H$>$A_H$, $T_H$>$G_H$, $C_H$>$A_H$ and $G_H$>$T_H$ - see the main text, chapter 2) into the initial multiple linear model. Next, following the logic of the stepwise backward multiple model (removing one the most non-significant variable at each step) we determined that three types of substitutions were independently associated with the generation length (*GL*).

*log2(GL) ~ 10.39 + 0.26*($A_H$>$G_H$) - 0.18*($T_H$>$A_H$) - 0.16*($C_H$>$A_H$),* equation (I)
*N = 424, $R^2$=0.104,*
*total p-value = 1.249e-10; p-values of transversions are < 0.002, p-value of $A_H$>$G_H$ = 6.42e-06,*
*presented coefficients are scaled.*

Comparing the scaled regression coefficients we observed that the strongest (the highest absolute effect size), as well as the most significant (the lowest p-value) effect was associated with $A_H$>$G_H$ transitions. Positive correlation of generation length with the frequency of a $A_H$>$G_H$ transition and negative correlation with the frequencies of $C_H$>$A_H$ and $T_H$>$A_H$ transversions as expected lead to the increased Ts/Tv in long-lived species described above.

It is important to note that the inclusion into the linear model (equation I) of the total number of mutations used to estimate the species-specific mutational spectrum (synonymous mutations in four-fold degenerate sites in *MT-CYB* gene) does not affect these results quantitatively:

*log2(GL) ~ 10.39 + 0.24*($A_H$>$G_H$) - 0.14*($T_H$>$A_H$) - 0.17*($C_H$>$A_H$) - 0.26*(Number Of Mutations)* equation (Ia)
*N = 424, $R^2$=0.145,*
*total p-value = 2.464e-14; p-value ($A_H$>$G_H$) = 1.66e-05; p-value ($T_H$>$A_H$) = 0.0103; p-value ($C_H$>$A_H$) = 0.00249; p-value (Number Of Mutations) = 5.29e-06; presented coefficients are scaled.*

We can see that the number of mutations negatively correlates with GL, probably because short-lived, small-bodied mammals are better investigated and have more sequences deposited in GenBank. Importantly, all effects demonstrated in the main text and in the equation I stay qualitatively similar.

Similarly with PGLS analyses performed before (supplementary materials 2.3) we repeated three models, analyzing an association of $A_H$>$G_H$ with the generation length. The final model 2.4C (table S2C) shows a significant positive effect of the generation length on the frequency of $A_H$>$G_H$ and the absence of the effect of the 'number of mutations'

*Table S2C. Phylogenetic generalised least-squares regression (PGLS) models, describing the relationships between the substitution rate $A_H$>$G_H$ and the generation length.*

| ***model*** | ***variable*** | ***coefficients*** | ***p values*** |
|---|---|---|---|
| ***2.4A: $A_H$>$G_H$ ~ log2(generation length)**; N=211, lambda [ ML] = 0.336* | *intercept* | *0.0202212* | *0.8184* |
| | *log2(generation length)* | *0.0115248* | *0.1488* |
| ***2.4B: $A_H$>$G_H$ ~ 0 + log2(generation length)**; N=211, lambda [ ML] = 0.317* | *log2(generation length)* | *0.0133441* | *3.173e-08* |
| ***2.4C: $A_H$>$G_H$ ~ 0 + log2(generation length) + log2(number of mutations)**; N=211, lambda [ ML] = 0.329* | *log2(generation length)* | *0.0123511* | *0.0003097* |
| | *log2(number of mutations)* | *0.0018587* | *0.6942430* |

(2.5) The principal component analysis (PCA): the second PC represents a signature of a longevity-associated mutagen (https://github.com/polarsong/mtDNA_mutspectrum/tree/VertebratePolymorphisms/Head/2Scripts/*VertebratePolymorphisms.MutSpecComparisons.Analyses.Ecology.Mammals.R*):

The correlation of the second principal component with the generation length was significantly stronger (rho = -0.39, p < 2.2e-16, N = 424) as compared to the sole effect of $A_H$>$G_H$ (Spearman's rho = 0.252, p value = 1.188e-07, N = 424) and the sole effect of Ts/Tv (Spearman's rho = 0.23, p-value = 2.021e-06; N = 424), suggesting that this second component could reflex a complex signature of a specific mutagen associated with the generation length. Indeed, both transversions $T_H$>$A_H$ and $C_H$>$A_H$, negatively associated with generation length (equation I), have strong effects on the second principal component and, as expected, point to the opposite direction compared to $A_H$>$G_H$ (Figure 2C left panel). Analysis of a subset of species with many (more than 60) mutations, used to reconstruct the mutational spectrum, demonstrated quantitatively similar results (data not shown).

To investigate if the second principal component is affected by the total number of mutations, used to reconstruct the mutational spectrum, we performed several analyses. We demonstrated that the second principal component positively correlated with the number of mutations (Spearman's rho = 0.2247093, p = 2.964e-06). However, in the multiple linear model where the second principal component was a function of both generation length and the number of mutations the effect of generation length is stronger and more significant:

PCA2 ~ -0.34*GL + 0.16*(Number Of Mutations); *equation (Ib)*
*N = 424, $R^2$=0.109,*
*total p-value* = 1.121e-11*; p-value (GL)* = 7.07e-09*; p-value (Number Of Mutations)* = 0.006*;*
*presented coefficients are scaled.*

The first principal component, loaded mainly by the most common $C_H$>$T_H$ transition, appears to represent an unknown source of variation, which we were unable to associate with any life-history traits such as generation length, body temperature, body mass, metabolic rate or the number of mutations used to derive the mutational spectrum.

<u>(2.6) The ratio of the two most common transitions also demonstrates an association with the generation length:</u>

*github:*
[https://github.com/polarsong/mtDNA_mutspectrum/tree/VertebratePolymorphisms/Head/2Scripts/*VertebratePolymorphisms.MutSpecComparisons.Analyses.Ecology.Mammals.R*](https://github.com/polarsong/mtDNA_mutspectrum/tree/VertebratePolymorphisms/Head/2Scripts/VertebratePolymorphisms.MutSpecComparisons.Analyses.Ecology.Mammals.R)

Since low frequency genetic variants might occur due to DNA damage during sequencing (Chen et al. 2017, 2018; Stewart et al. 2018) we replicated our results taking into account only two the most common transitions: $A_H$>$G_H$ and $C_H$>$T_H$. Analysing the fraction of $A_H$>$G_H$ (derived as $A_H$>$G_H$ / ($A_H$>$G_H$ + C>T)), we still observed the positive correlation of this fraction with generation length (Spearman's rho = 0.21, p = 8.441e-06, n = 424).

Similarly with PGLS analyses performed before (supplementary materials 2.3) we repeated three models, analyzing an association of $A_H$>$G_H$ / ($A_H$>$G_H$ + C>T) with the generation length. The final model 2.6C (table S2D) shows a significant positive effect of the generation length on $A_H$>$G_H$ / ($A_H$>$G_H$ + C>T) and the absence of the effect of the 'number of mutations'.

*Table S2D. Phylogenetic generalised least-squares regression (PGLS) models, describing the relationships between $A_H$>$G_H$ / ($A_H$>$G_H$+ $C_H$>$T_H$) and generation length.*

| ***model*** | ***variable*** | ***coefficients*** | ***p values*** |
|---|---|---|---|
| ***2.6A: $A_H$>$G_H$ / ($A_H$>$G_H$+ $C_H$>$T_H$) ~ log2(generation length)****;* *N=211, lambda [ ML] = 0.000* | *intercept* | *-0.0353585* | *0.6318952* |
| | *log2(generation length)* | *0.0239113* | *0.0005575* |
| ***2.6B: $A_H$>$G_H$ / ($A_H$>$G_H$+ $C_H$>$T_H$) ~ 0 + log2(generation length)****;* *N=211, lambda [ ML] = 0.000* | *log2(generation length)* | *0.0206577* | *< 2.2e-16* |
| ***2.6C:$A_H$>$G_H$ / ($A_H$>$G_H$+ $C_H$>$T_H$) ~ 0 + log2(generation length) + log2(number of mutations)****;* *N=211, lambda [ ML] = 0.000* | *log2(generation length)* | *0.0225146* | *6.665e-12* |
| | *log2(number of mutations)* | *-0.0034229* | *0.5373* |

<u>(2.7) A nucleotide-content-independent mutational spectrum demonstrates a positive correlation between the generation length and $A_H$>$G_H$:</u>

*github:*
[https://github.com/polarsong/mtDNA_mutspectrum/tree/VertebratePolymorphisms/Head/2Scripts/*VertebratePolymorphisms.MutSpecComparisons.Analyses.Ecology.Mammals.R*](https://github.com/polarsong/mtDNA_mutspectrum/tree/VertebratePolymorphisms/Head/2Scripts/VertebratePolymorphisms.MutSpecComparisons.Analyses.Ecology.Mammals.R)

The fraction of $A_H$>$G_H$ in each species-specific mutational spectra depends on both the number of observed $A_H$>$G_H$ substitutions within a species and the number of $A_H$ nucleotides in fourfold degenerate synonymous position of *MT-CYB* of a given species, used for normalization (see supplementary materials 2.2). To demonstrate that the observed above results were not solely driven by the variation in the frequency of ancestral nucleotides (i.e. by denominator in the process of normalization), we recalculated $A_H$>$G_H$ fraction for each species using only substitutions from $A_H$: $A_H$>$G_H$ / ($A_H$>$G_H$ + $A_H$>$C_H$ + $A_H$>$T_H$). Using this approach we don't take into account nucleotide content of different species. We observed a positive correlation between $A_H$>$G_H$ and the mammalian generation length (Spearman's rho = 0.164, p = 0.0007, N = 424), suggesting that $A_H$>$G_H$ is higher in long-lived mammals irrespectively of the nucleotide content.

Similarly with PGLS analyses performed before (supplementary materials 2.3) we repeated models, analyzing an association of $A_H$>$G_H$ / ($A_H$>$G_H$+ $A_H$>$T_H$ + $A_H$>$C_H$) with generation length. The final model 2.7B (table S2E) shows a significant positive effect of the generation length on $A_H$>$G_H$ / ($A_H$>$G_H$+ $A_H$>$T_H$ + $A_H$>$C_H$) and the absence of the effect of the 'number of mutations'.

*Table S2E. Phylogenetic generalised least-squares regression (PGLS) models, describing the relationships between $A_H$>$G_H$ / ($A_H$>$G_H$+ $A_H$>$T_H$ + $A_H$>$C_H$) and generation length.*

| ***model*** | ***variable*** | ***coefficients*** | ***p values*** |
|---|---|---|---|
| ***2.7A: $A_H$>$G_H$ / ($A_H$>$G_H$+ $A_H$>$T_H$ + $A_H$>$C_H$) ~ log2(generation length)****;* *N=211, lambda [ ML] = 0.000* | *intercept* | *0.6214988* | *1.759e-12* |
| | *log2(generation length)* | *0.0208230* | *0.007154* |
| ***2.7B: $A_H$>$G_H$ / ($A_H$>$G_H$+ $A_H$>$T_H$ + $A_H$>$C_H$) ~ log2(generation length) + log2(number of mutations)****;* *N=211, lambda [ ML] = 0.000* | *intercept* | *0.6581961* | *1.655e-08* |
| | *log2(generation length)* | *0.0196741* | *0.01515* |
| | *log2(number of mutations)* | *-0.0041067* | *0.62634* |

**(3) mtDNA of mammals with high generation length are more Ah poor and Gh rich due to intensive Ah>Gh mutagenesis**

(3.1) Mammals are close to their compositional nucleotide equilibrium:

*github:*
*https://github.com/polarsong/mtDNA_mutspectrum/blob/WholeGenomesBranch/Head/2Scripts/EquilibriumAnalysMammals.R*

*Table S3A. An average mutational spectra of mammals with very short and very long generation length was used as input to simulations. As expected,* $A_H$>$G_H$ *is almost two times higher in species with very long versus very short generation length.*

| Average mutational spectrum of mammals with very short generation length (less than the lower decile: 554 days, N = 27) | | Average mutational spectrum of mammals with very long generation length (higher than the upper decile: 5221 days, N =25) | |
|---|---|---|---|
| $T_H$>$A_H$ | 0.0060787 | $T_H$>$A_H$ | 0.0026855 |
| $T_H$>$C_H$ | 0.0174578 | $T_H$>$C_H$ | 0.0399171 |
| $T_H$>$G_H$ | 0.0049765 | $T_H$>$G_H$ | 0.0030254 |
| $A_H$>$T_H$ | 0.0191651 | $A_H$>$T_H$ | 0.0123239 |
| $A_H$>$C_H$ | 0.0028298 | $A_H$>$C_H$ | 0.0031279 |
| **$A_H$>$G_H$** | **0.1239227** | **$A_H$>$G_H$** | **0.2173696** |
| $C_H$>$T_H$ | 0.6664678 | $C_H$>$T_H$ | 0.5693657 |
| $C_H$>$A_H$ | 0.0392862 | $C_H$>$A_H$ | 0.0317306 |
| $C_H$>$G_H$ | 0.0189419 | $C_H$>$G_H$ | 0.0132412 |
| $G_H$>$T_H$ | 0.0155279 | $G_H$>$T_H$ | 0.0169743 |
| $G_H$>$A_H$ | 0.0832090 | $G_H$>$A_H$ | 0.0871418 |
| $G_H$>$C_H$ | 0.0021366 | $G_H$>$C_H$ | 0.0030971 |

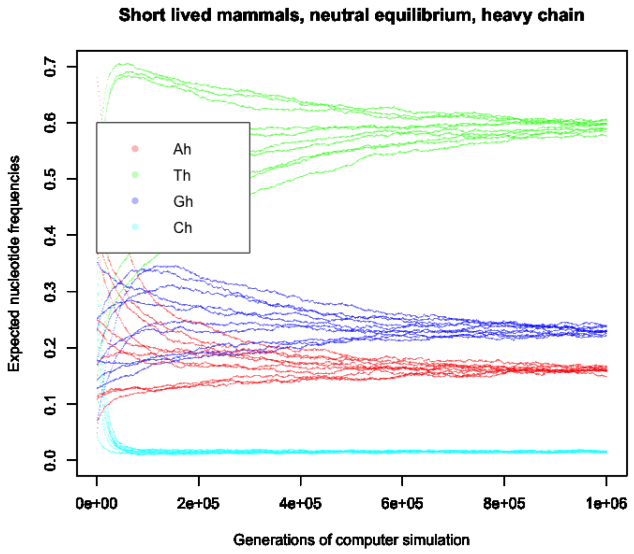


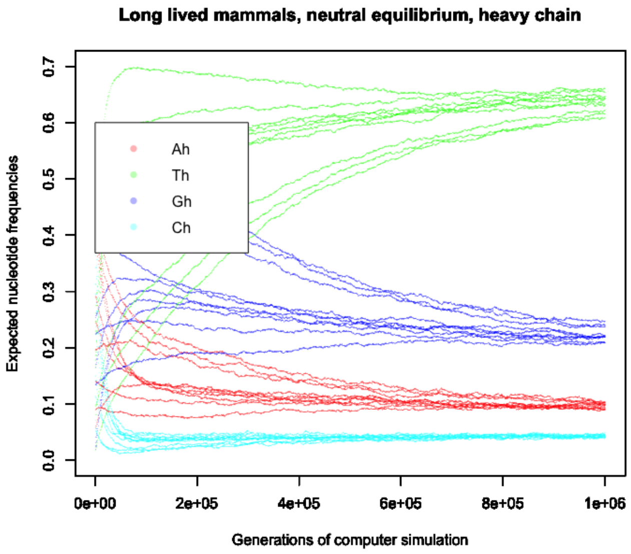

*Figure S3A* Expected nucleotide composition of mammals, obtained as a result of simulations. We can see that species with higher generation length (right panel) are characterized by the increased frequency of Gh and decreased frequency of Ah. An expected nucleotide content at compositional equilibrium was derived using R simulation. The main goal of the simulation was to obtain an equilibrium nucleotide composition (frequencies of four nucleotides) evolved under a given mutational spectrum. Simulations started with random frequencies of four nucleotides (left part of the plots) and with time (with the simulated generations) the nucleotide frequencies changed according to the mutational spectrum, reaching the saturation (right part of the plots). Curves corresponding to the nucleotide frequencies converge during the simulation to one value. These equilibrium values (4 nucleotide frequencies at the end of the simulation) were used in our downstream analyses as expected nucleotide content (see Fig 3A in the main text).

(3.2) The mutational bias affects neutral nucleotide composition of complete mammalian mitochondrial genomes:

Table S3B. *Ah>Gh frequency in polymorphism-derived 12-component mutational spectrum affects the nucleotide content in four-fold degenerate synonymous positions. As expected, high Ah>Gh substitution rate increases GhAh skew, increases the frequency of Gh and decreases the frequency of Ah.*

| ***class*** | ***method*** | ***variable*** | ***Spearman's Rho*** | ***P value*** |
|---|---|---|---|---|
| **Mammalia (N = 247)** | **pairwise spearman's correlation** | **GhAh Skew** | **0.486147** | **4.677e-16** |
| | | **Fraction Ah** | **-0.4913123** | **< 2.2e-16** |
| | | **Fraction Gh** | **0.409863** | **1.824e-11** |

*github:*
*https://github.com/polarsong/mtDNA_mutspectrum/blob/WholeGenomesBranch/Body/3Results/AllGenesCodonUsageNoOverlap.txt*

*https://github.com/polarsong/mtDNA_mutspectrum/blob/WholeGenomesBranch/Head/2Scripts/WholeGenomeAnalyses.NoOverlap.AGSkewAndGradient.R*

Effect of the mutational spectrum on neutral nucleotide content: high $A_H$>$G_H$ in long-lived species leads to a decrease in $A_H$ and increase in $G_H$ as shown by rank correlations (Table S3C), stepwise backward multiple linear models (Table S3C) and hylogenetically aware statistics (see below).

*Table S3C. Correlations between generation time and nucleotide fractions in neutral sites*

| **class** | **method** | **nucleotide** | **Spearman's Rho / coefficient in linear model** | **P value** |
|---|---|---|---|---|
| Mammalia (N = 650) | pairwise spearman's correlation | $T_H$ | -0.27 | 3.635e-12 |
| | | $A_H$ | -0.31 | 1.287e-15 |
| | | $C_H$ | 0.18 | 3.665e-06 |
| | | $G_H$ | 0.47 | < 2.2e-16 |

| | | | | |
|---|---|---|---|---|
| | results of the backward stepwise multiple linear model | intercept | 11.06 | < 2.2e-16 |
| | | $A_H$ | -0.11 | 0.023 |
| | | $G_H$ | 0.46 | < 2.2e-16 |

Consideration of phylogenetic inertia by means of phylogenetically independent contrasts demonstrated the same trend (fraction of $A_H$: spearman's rho = - 0.09, p = 0.025;  fraction of $G_H$: spearman's rho = 0.09, p = 0.021).

(3.3) Asymmetry of codon usage driven by two the most common transitions as a function of generation length

*github:*
https://github.com/polarsong/mtDNA_mutspectrum/blob/WholeGenomesBranch/Head/2Scripts/AsymmetryCodonUsage.R

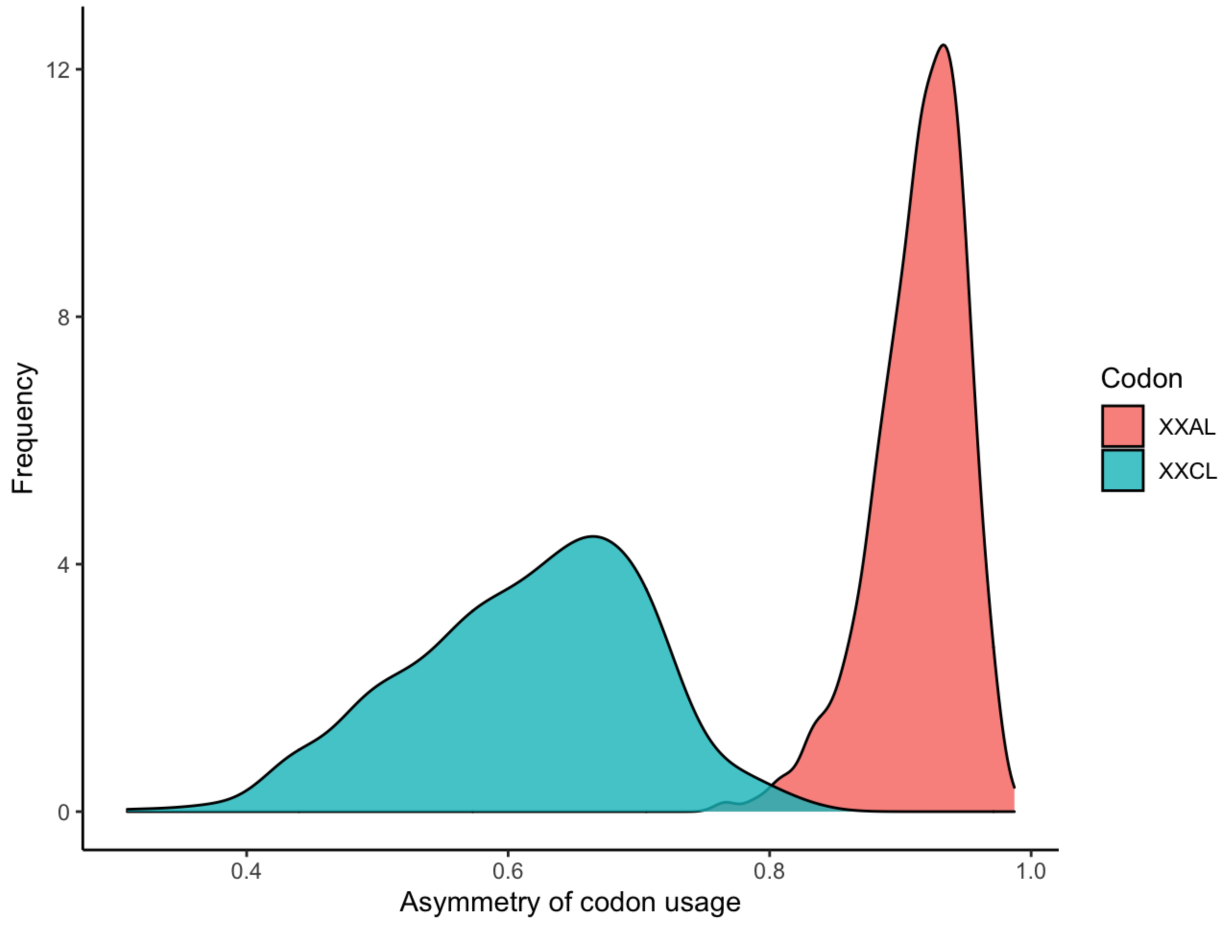


*Figure S3B Distribution of asymmetry of $XXA_L$ and $XXC_L$ codon usage. Mann-Whitny paired U test (N = 648) showed more pronounced asymmetry of $XXA_L$ as compared to $XXC_L$ (p-value < 2.2e-16), while both of them significantly deviate from expected 0.5 (p-values are  < 2.2e-16, Wilcoxon test with mu = 0.5).*

*Table S3D. Rank correlations of the codon asymmetries with the generation length.*

| class | method | Codon asymmetry | Spearman's Rho | P value |
|---|---|---|---|---|
| Mammalia (N = 648) | pairwise spearman's correlation | $XXC_L$ | 0.3437 | 2.2e-16 |
| | | $XXA_L$ | -0.2233 | 9.164e-09 |

*Table S3E standard linear models and PGLS models of the association of the codon asymmetries with the generation length. Only $XXC_L$ shows robust association with the generation length (see models 3.3C and 3.3D).*

| ***model*** | ***variable*** | ***coefficients*** | ***p values*** |
|---|---|---|---|
| **3.3A: $XXA_L$~log2(generation length)**<br>*N=648* | *intercept* | *0.285261* | *<2e-16* |
| | *log2(generation length)* | *0.029795* | *<2e-16* |
| ***3.3B: $XXC_L$ ~ log2(generation length)***<br>*N=648* | *intercept* | *0.993573* | *<2e-16* |
| | *log2(generation length)* | *-0.007203* | *5.15e-09* |
| **3.3C: scale($XXA_L$)~log2(generation length)**<br>*N=648, lambda [ ML] = 0.998* | *intercept* | *1.001718* | *0.27865* |
| | *log2(generation length)* | *-0.084755* | *0.09565* |
| ***3.3D: scale($XXC_L$) ~ log2(generation length)***<br>*N=648; lambda [ ML] = 1.000* | *intercept* | *-1.416211* | *0.03242* |
| | *log2(generation length)* | *0.076297* | *0.02704* |

**(4) nucleotide composition is a function of both time spent single stranded (TSSS) and the generation length**

*(4.1) $G_H A_H$ skew is a function of both TSSS and GT*

(github:https://github.com/polarsong/mtDNA_mutspectrum/blob/WholeGenomesBranch/Head/2Scripts/WholeGenomeAnalyses.NoOverlap.AGSkewAndGradient.R)

Having two factors, the time spent single stranded (TSSS) and the generation length (GL), affecting nucleotide composition of mtDNA we first, analysed how $G_H A_H$ skew depends on both TSSS and GL (Table S4A). We observed that both factors affect the skew significantly and an interaction term (see model 4C) was non significant.

Models 4B (with scaled variables) and 4D (with Dummy variables) allow us to compare effect sizes of the TSSS and GL: how strongly they affect the $G_H A_H$ skew? Although GL coefficient is less than the TSSS coefficient they are comparable, suggesting that generation length is as important as the position of the gene along the major arc.

Analyses, performed separately for two the biggest in our dataset families (Laurasiatheria and *Ruminantia*) and their corresponding models 4F, 4H, 4J, 4L confirmed the importance of both these factors (TSSS and GL) on $G_H A_H$ skew.

| ***model*** | ***variable*** | ***coefficients*** | ***p values*** |
|---|---|---|---|
| **4A:** $G_H A_H$ Skew~ TSSS + log2(generation length) *N=648* | *intercept* | -0.8498809 | *<2e-16* |
| | *log2(generation length)* | *0.0724620* | *<2e-16* |
| | *TSSS* | *0.0387322* | *<2e-16* |
| **4B:** $G_H A_H$ Skew~ scale(TSSS) + scale(generation length) *N=648* | *intercept* | *0.164223* | *<2e-16* |
| | *scale(generation length)* | *0.069142* | *<2e-16* |
| | *scale(TSSS)* | *0.111258* | *<2e-16* |
| **4C:** $G_H A_H$Skew ~ TSSS*log2(generation length) *N=648* | *intercept* | *-0.8619907* | *<2e-16* |
| | *log2(generation length)* | *0.0735574* | *<2e-16* |
| | *TSSS* | *0.0409340* | *2.66e-05* |
| | *log2(generation length):TSSS* | *-0.0001992* | *0.82* |
| **4D:** $G_H A_H$Skew ~ *DummyHighTsss + DummyLonglived* *N=648* | *intercept* | *0.0002864* | *0.955* |
| | *DummyLonglived* | *0.1385817* | *<2e-16* |
| | *DummyHighTSSS* | *0.1903651* | *<2e-16* |

*Table S4A. $G_H A_H$ skew is a positive function of both TSSS and GT. The analyzed sample size is all mammalian species (N = 648). Dummy variables (DummyLonglived equals 1 for mammals with GT higher than median and zero, otherwise; DummyHighTSSS equals 1 for genes COX1, COX2, ATP6, ATP8, COX3 and zero for ND3, ND4L, ND5, CytB) were introduced into the models to show robustness of the associations.*

| order | model | variable | coefficients | p values |
| --- | --- | --- | --- | --- |
| **Laurasiatheria (N=*126*)** | **4E:** $G_HA_H$ Skew~ TSSS + log2(generation length) | *intercept* | -1.341080 | *<2e-16* |
| | | *log2(generation length)* | *0.109202* | *<2e-16* |
| | | *TSSS* | *0.042084* | *<2e-16* |
| | **4F:** $G_HA_H$ Skew~ scale(TSSS) + scale(generation length) | *intercept* | *0.084474* | *<2e-16* |
| | | *scale(generation length)* | *0.134692* | *<2e-16* |
| | | *scale(TSSS)* | *0.120924* | *<2e-16* |
| | **4G:** $G_HA_H$ Skew~ TSSS * log2(generation length) | *intercept* | *-1.346* | *<2e-16* |
| | | *log2(generation length)* | *1.096e-01* | *<2e-16* |
| | | *TSSS* | *4.292e-02* | *0.0239* |
| | | *log2(generation length) : TSSS* | *-7.688e-05* | *0.9644* |
| | **4H:** $G_HA_H$ Skew~ *DummyHighTSSS* + *DummyLonglived* | *intercept* | *-0.13240* | *<2e-16* |
| | | *DummyLonglived* | *0.25064* | *<2e-16* |
| | | *DummyHighTSSS* | *0.20299* | *<2e-16* |
| ***Ruminantia* (N=*112)*** | **4I:** $G_HA_H$ Skew~ TSSS + log2(generation length) | *intercept* | -0.577119 | *0.000431* |
| | | *log2(generation length)* | *0.047036* | *0.001197* |
| | | *TSSS* | *0.043887* | *<2e-16* |
| | **4J:** $G_HA_H$ Skew~ scale(TSSS) + scale(generation length) | *intercept* | *0.193266* | *<2e-16* |
| | | *scale(generation length)* | *0.023978* | *0.000159* |
| | | *scale(TSSS)* | *0.126113* | *<2e-16* |
| | **4K:** $G_HA_H$ Skew~ TSSS * log2(generation length) | *intercept* | *-1.105786* | *0.00172* |
| | | *log2(generation length)* | *0.094042* | *0.00268* |
| | | *TSSS* | *0.140008* | *0.01369* |
| | | *log2(generation length):TSSS* | *-0.008547* | *0.09006* |

| | | | | |
|---|---|---|---|---|
| | **4L:** $G_HA_H$Skew ~ *DummyHighTss + DummyLonglived* | *intercept* | *0.05074* | *<2e-16* |
| | | *DummyLonglived* | *0.06689* | *1.99e-06* |
| | | *DummyHighTSSS* | *0.19905* | *<2e-16* |

Table S4B. *$G_HA_H$ skew shows a positive associations with both TSSS and GT within two the biggest mammalian families: Laurasiatheria and Ruminantia. Dummy variables (DummyLonglived equals 1 for mammals with GT higher than median and zero, otherwise; DummyHighTSSS equals 1 for genes COX1, COX2, ATP6, ATP8, COX3 and zero for ND3, ND4L, ND5, CytB) were introduced into the models to show robustness of the associations.*

*(4.2) Nucleotide gradient (decrease in $A_H$ and increase in $G_H$) along mtDNA tend to be more pronounced in species with high generation length*

We were interested if the slopes of changes in nucleotide content (Figure 4) differed between short- and long-lived species. We plotted nucleotide content at neutral sites of each of 13 genes and correlated it with gene location, ranked from the shortest (MT-CO1) to the longest time (MT-CYB) of being single-stranded (Figure S4). We observed that the frequency of $G_H$ increased along the gradient of time of being single stranded (Spearman's Rho >= 0.62, p-values <= 0.028). Given the fact that this analyses was limited by the low number of genes (13), we performed more sensitive analysis splitting the genome of each species into 50 windows containing 25 neutral (fourfold degenerate synonymous) nucleotides and using only genes located on the same strand of the major arc (all genes except ND1, ND2 and ND6). For each window we estimated the frequencies of four nucleotides and for each species we calculated the Spearman Rho coefficient and p-value, estimating the changes in a given metric along the genome (Figure S4). We can see that the frequency of $G_H$ indeed has the strongest bias towards positive rho values, and additionally we can see also, that the second strongest bias belongs to $A_H$, which decreased along the genome (Figure S4).

To test additionally if some of these gradients are more pronounced in long- versus short-lived mammals, we estimated the fraction of long-lived mammals (with generation length more than median, 1101.1 day) among significant (with p-values <=0.01) rho values. Only for $G_H$ nucleotides we observed significant excess of long-lived mammals among the significant ones (Figure S4, odds ratio = 1.61, p-value = 0.003521 for $G_H$ nucleotide, Fisher test). This suggests that the rate of increase in the frequency of $G_H$ is faster in long- versus short- lived mammals.

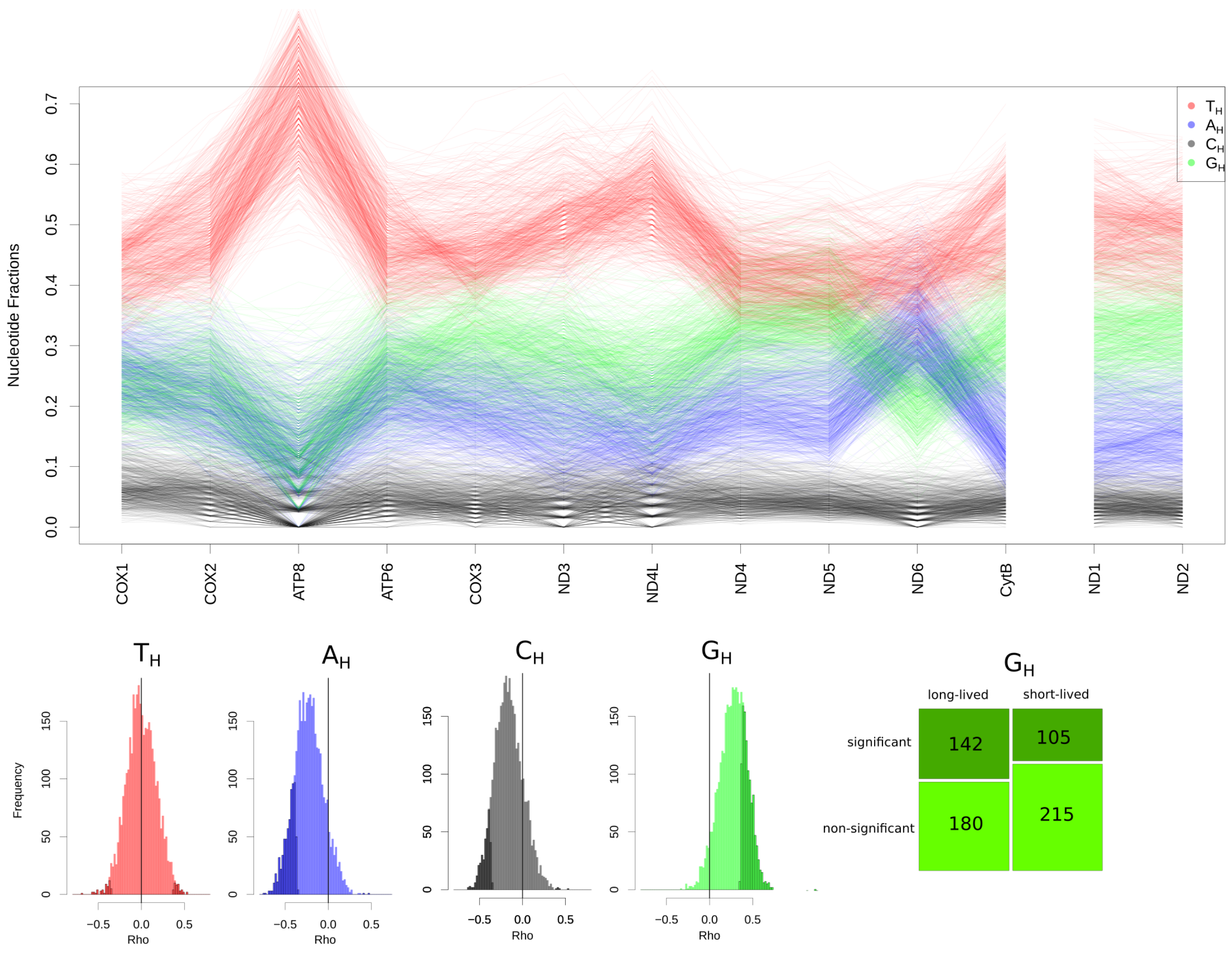


*Figure S4. The long-term effect of the mutational bias: neutral nucleotide content in mammalian species*

*Upper panel. Changes in nucleotide content along mammalian mtDNA. (N = 650). All genes located in the major arc are ranked according to the time spent single stranded: from COX1 to CYTB. ND2 gene spent more time than ND1 in the single-strand state but we do not compare directly these two genes with all others from the major arc.*

*Lower panel. The gradient of nucleotide changes with time being single stranded (increase in $G_H$) is more pronounced in long-lived mammals (N = 650). Histograms of spearman Rho values demonstrate that $A_H$ is decreasing with the time being single stranded while $G_H$ is increasing.*

*Mosaicplot reflects an excess of long-lived mammals among species with significant Rho for $G_H$, reflecting the higher* rate of increase in the frequency of $G_H$ *in mtDNA of long-lived mammals.*

**(5) The frequencies of $A_H$ ($T_L$) and $G_H$ ($C_L$) at the third, the most neutral, codon position ($XXT_L$ and $XXC_L$) demonstrate the strongest associations with the generation length.**

Out of 12 regressions (4 types of nucleotides x 3 positions in a codon) of the nucleotide composition against the generation length only 4 regressions with $T_L$ and $C_L$ demonstrate significant effects ($T_LXX$, $XXT_L$, $C_LXX$, $XXC_L$ , X marks any nucleotide, all p -values < 0.01 are marked as a bold font in the table). The signs of the regression coefficients (negative for $T_L$ and positive for $C_L$) are in line with more intensive $A_H$>$G_H$ ($T_L$>$C_L$) mutagenesis in mammals with high generation length. The strongest effect size (an absolute value of the regression coefficient) is observed for the third, the most neutral codon position (-16.1202 for $XXT_L$ and 16.8590 for $XXC_L$), which is about four times higher as compared to the effect size of the first codon position (-3.7802 in case of $T_LXX$ and 4.6674 for $C_LXX$). Altogether these results support our main result that the $A_H$>$G_H$ ($T_L$>$C_L$) mutagenesis is the strongest factor shaping an association of the mtDNA nucleotide content with the generation length. The observation that the first codon positions show significant, although low-effect size, associations suggests that selection can also play some role. More studies are needed to dissect it.

All results were obtained using phylogenetic generalized linear models (PGLS). In several cases PGLS models didn't converge (marked as '-' in the table).

(https://github.com/polarsong/mtDNA_mutspectrum/blob/WholeGenomesBranch/Head/2Scripts/Codon_Comp.R)

| ***Model*** <br> ***(X marks any nucleotide in the codon)*** | ***Variable*** | ***Coefficients*** | ***P values (nominal)*** |
|---|---|---|---|
| **$T_L$XX**~ log2(GenerationLength) <br> PGLS, lambda [ML] : 1.000, <br> N = 648 | intercept | 881.6297 | < 2.2e-16 *** |
| | log2(GenerationLength) | **-3.7802** | **0.008283 **** |
| X$T_L$X~ log2(GenerationLength) <br> PGLS, lambda [ML] : 0.999, <br> N = 648 | intercept | 1517.11145 | < 2e-16 *** |
| | log2(GenerationLength) | -1.17336 | 0.05864 |
| **XX$T_L$** ~ log2(GenerationLength) <br> PGLS, lambda [ML] : 1.000, <br> N = 648 | intercept | 1077.1516 | < 2.2e-16 *** |
| | log2(GenerationLength) | **-16.1202** | **0.003075 **** |
| **$C_L$XX** ~ log2(GenerationLength) <br> PGLS, lambda [ML] : 1.000, <br> N = 648 | intercept | 832.3117 | < 2.2e-16 *** |
| | log2(GenerationLength) | **4.6674** | **0.001571 **** |
| X$C_L$X ~ log2(GenerationLength) <br> PGLS, lambda [ML] : 1.000, <br> N = 648 | intercept | 943.39369 | <2e-16 *** |
| | log2(GenerationLength) | 0.72420 | 0.2196 |
| **XX$C_L$** ~ log2(GenerationLength) <br> PGLS, lambda [ML] : 1.000, <br> N = 648 | intercept | 856.8488 | 3.109e-15 *** |
| | log2(GenerationLength) | **16.8590** | **0.002354 **** |
| $A_L$XX ~ log2(GenerationLength) <br> PGLS, lambda [ML] : 1.000, <br> N = 648 | intercept | 1134.12953 | < 2e-16 *** |
| | log2(GenerationLength) | -1.43611 | 0.08783 |
| X$A_L$X ~ log2(GenerationLength) <br> PGLS, lambda [ML] : 0.119, <br> N = 648 | intercept | **-** | **-** |
| | log2(GenerationLength) | **-** | **-** |

| | | | |
|---|---|---|---|
| $XXA_L$~ log2(GenerationLength) PGLS, lambda [ML] : 1.000, N = 648 | intercept | 1514.3797 | <2e-16 *** |
| | log2(GenerationLength) | -3.9375 | 0.262 |
| $GXX_L$~ log2(GenerationLength) PGLS, lambda [ML] : 1.000, N = 648 | intercept | 723.20809 | <2e-16 *** |
| | log2(GenerationLength) | 0.68135 | 0.3585 |
| $XGX_L$~ log2(GenerationLength) PGLS, lambda [ML] : 0.119, N = 648 | intercept | - | - |
| | log2(GenerationLength) | - | - |
| $XXG_L$~ log2(GenerationLength) PGLS, lambda [ML] : 1.000, N = 648 | intercept | 122.8989 | 0.01092 * |
| | log2(GenerationLength) | 3.3311 | 0.18482 |

**References**


"Bionumbers." n.d. Accessed December 24, 2020. http://book.bionumbers.org/how-quickly-do-different-cells-in-the-body-replace-themselves/.

Chen, Lixin, Pingfang Liu, Thomas C. Evans Jr, and Laurence M. Ettwiller. 2017. "DNA Damage Is a Pervasive Cause of Sequencing Errors, Directly Confounding Variant Identification." *Science* 355 (6326): 752–56.

———. 2018. "Response to Comment on 'DNA Damage Is a Pervasive Cause of Sequencing Errors, Directly Confounding Variant Identification.'" *Science*. https://doi.org/10.1126/science.aat0958.

Kim, Jung Yeon, Simon Tavaré, and Darryl Shibata. 2005. "Counting Human Somatic Cell Replications: Methylation Mirrors Endometrial Stem Cell Divisions." *Proceedings of the National Academy of Sciences of the United States of America* 102 (49): 17739–44.

Rebolledo-Jaramillo, B., M. S. -W. Su, N. Stoler, J. A. McElhoe, B. Dickins, D. Blankenberg, T. S. Korneliussen, et al. 2014. "Maternal Age Effect and Severe Germ-Line Bottleneck in the Inheritance of Human Mitochondrial DNA." *Proceedings of the National Academy of Sciences* 111 (43): 15474–79.

Seim, Inge, Siming Ma, and Vadim N. Gladyshev. 2016. "Gene Expression Signatures of Human Cell and Tissue Longevity." *Npj Aging and Mechanisms of Disease* 2 (1): 1–8.

Stewart, Chip, Ignaty Leshchiner, Julian Hess, and Gad Getz. 2018. "Comment on 'DNA Damage Is a Pervasive Cause of Sequencing Errors, Directly Confounding Variant Identification.'" *Science*. https://doi.org/10.1126/science.aas9824.

Tomasetti, Cristian, Rick Durrett, Marek Kimmel, Amaury Lambert, Giovanni Parmigiani, Ann Zauber, and Bert Vogelstein. 2017. "Role of Stem-Cell Divisions in Cancer Risk." *Nature* 548 (7666): E13–14.

Tomasetti, Cristian, and Bert Vogelstein. 2015. "Cancer Etiology. Variation in Cancer Risk among Tissues Can Be Explained by the Number of Stem Cell Divisions." *Science* 347 (6217): 78–81.

"Urothelial Stem Cell Regeneration." n.d. Accessed December 24, 2020. https://reproductivesciences.wustl.edu/laboratories/mysorekar-lab/urothelial-stem-cell-regeneration/.

Wang, Caihong, Whitney Trotter Ross, and Indira U. Mysorekar. 2017. "Urothelial Generation and Regeneration in Development, Injury, and Cancer." *Developmental Dynamics: An Official Publication of the American Association of Anatomists* 246 (4): 336.

Wei, Wei, Salih Tuna, Michael J. Keogh, Katherine R. Smith, Timothy J. Aitman, Phil L. Beales, David L. Bennett, et al. 2019. "Germline Selection Shapes Human Mitochondrial DNA Diversity." *Science* 364 (6442). https://doi.org/10.1126/science.aau6520.

Yue Li, Rebecca A. Wingert. 2013. "Regenerative Medicine for the Kidney: Stem Cell Prospects & Challenges." *Clinical and Translational Medicine* 2: 11.

Zaidi, Arslan A., Peter R. Wilton, Marcia Shu-Wei Su, Ian M. Paul, Barbara Arbeithuber, Kate Anthony, Anton Nekrutenko, Rasmus Nielsen, and Kateryna D. Makova. 2019. "Bottleneck and Selection in the Germline and Maternal Age Influence Transmission of Mitochondrial DNA in Human Pedigrees." *Proceedings of the National Academy of Sciences of the United States of America* 116 (50): 25172–78.
